# Lipid Transfer Proteins and PI4KIIα Initiate Nuclear p53-Phosphoinositide Signaling

**DOI:** 10.1101/2023.05.08.539894

**Authors:** Noah D. Carrillo, Mo Chen, Tianmu Wen, Poorwa Awasthi, Colin Sterling, Dhruv Brahmbhatt, Vincent L. Cryns, Richard A. Anderson

## Abstract

Phosphoinositide (PIP_n_) messengers are present in non-membranous regions of nuclei where they are assembled into a phosphatidylinositol (PI) 3-kinase (PI3K)/Akt pathway that is distinct from the cytosolic membrane-localized pathway. In the nuclear pathway, PI kinases/phosphatases bind the p53 tumor suppressor protein (wild-type and mutant) to generate p53-PIP_n_ complexes (p53-PIP_n_ signalosome) that activate Akt by a PI3,4,5P_3_-dependent mechanism in non-membranous regions of the nucleus. This pathway is dependent on a source of nuclear PIP_n_s that is poorly characterized. Here we report that a subset of PI transfer proteins (PITPs), which transport PI between membranes to enable membrane-localized PIP_n_ synthesis, also interact with p53 in the nucleus upon genotoxic stress. Class I PITPs (PITPα/β) specifically supply the PI required for the generation of p53-PIP_n_ complexes and subsequent signaling in the nucleus. Additionally, the PI 4-kinase PI4KIIα binds to p53 and the PITPs to catalyze the formation of p53-PI4P. p53-PI4P is then sequentially phosphorylated to synthesize p53-PIP_n_ complexes that regulate p53 stability, nuclear Akt activation and genotoxic stress resistance. In this way, PITPα/β and PI4KIIα bind p53 and collaborate to initiate p53-PIP_n_ signaling by mechanisms that require PI transfer by PITPα/β and the catalytic activity of PI4KIIα. Moreover, the identification of these critical upstream regulators of p53-PIP_n_ signaling point to PITPα/β and PI4KIIα as potential therapeutic targets in this pathway for diseases like cancer.

**In Breif:** Phosphatidylinositol transfer proteins and a PI 4-kinase initiate nuclear p53-PIP_n_ signaling in membrane-free regions.

## Introduction

Phosphoinositide (PIP_n_) lipids are synthesized from phosphatidylinositol (PI) in membranes by PIP-metabolizing enzymes, including kinases and phosphatases, to produce a second messenger network comprised of seven distinct isomers that play key roles in many cellular pathways^1^. Cytosolic class I PI 3-kinases (PI3Ks) catalyze the synthesis of PI3,4,5P_3_, which leads to the recruitment and activation of the serine/threonine kinases phosphoinositide-dependent kinase 1 (PDK1), mTOR Complex 2 (mTORC2), and Akt at membranes^2–8^. Akt in turn phosphorylates many substrates, such as forkhead box O (FOXO) proteins, tuberous sclerosis complex 2 (TSC2), and BCL2-associated agonist of cell death (BAD), to promote cell growth and inhibit apoptosis^4^. This membrane-localized PI3K/Akt pathway is often hyperactivated in cancer and class I PI3K inhibitors are currently in the clinic^9, 10^.

Akt is also activated in the nucleus^11–16^. Recent studies have identified a non-membranous nuclear PI3K/Akt pathway dependent on p53^17, 18^, a tumor suppressor frequently mutated in cancer, resulting in gain-of-function oncogenic acitivities^19, 20^. In this nuclear pathway, wild-type and mutant p53 are sequentially modified by PIP_n_s and their associated PIP kinases/phosphatases. The type I PI4P 5-kinase α (PIPKIα) directly interacts with p53 and generates the p53-PI4,5P_2_ complex (p53-PIP_n_ signalosome) in response to stress^18^. PI5P4Kβ has also been functionally linked to p53 as deletion of both genes is lethal in mice, but the mechanism is poorly defined^21^. The nuclear PI 3-kinase IPMK (inositol polyphosphate multikinase) acts on p53-PI4,5P_2_ to synthesize p53-PI3,4,5P_3_ complexes, which then recruit and activate Akt in the non-membranous nucleoplasm^17, 18^. The nuclear Akt pathway is insensitive to PI3K inhibitors in the clinic^17^, highlighting its therapeutic importance.

Although multiple PIP_n_s and PIP kinases/phosphatases are present in the nucleoplasm^16, 22–24^, their function and regulation are poorly characterized compared to PIP_n_ signaling at cytosolic membranes. Specifically, prior studies have demonstrated the synthesis of PI in the ER, which is transported from the ER to other membranes, and shuttled between membranes by a family of lipid transfer proteins, the phosphatidylinositol transfer proteins (PITPs)^25^. To date, research has focused on the cytosolic function of PITPs^26–28^. Intriguingly, PITPβ was suggested to interact with p53^29^, prompting us to investigate the functional role of PITPs in regulating the synthesis of p53-PIP_n_ complexes. Here we demonstrate that the highly homologous class I PITPs interact with p53 in the nucleus upon genotoxic stress and are specifically required to generate p53-PIP_n_ complexes. Additionally, a PI 4-kinase, PI4KIIα, binds to both p53 and the PITPs and is also required to initiate p53-PIP_n_ complex formation and nuclear p53-PIP_n_ signaling. PITPα/β and PI4KIIα collaborate to synthesize p53-PI4P complexes that are sequentially phosphorylated by PIP kinases or dephosphorylated by phosphatases to regulate p53 stability, nuclear Akt activation and stress resistance. These findings also implicate PITPα/β and PI4KIIα as therapeutic targets to regulate the nuclear p53-PIP_n_ pathway.

## Results

### p53 interacts with class I PITPs in the nucleus

The reported interaction of PITPβ and p53 in a proteomic screen^29^ suggests that PITPs may play a functional role in generating nuclear p53-PIP_n_ complexes. To explore this, we investigated the interaction of p53 with all five PITPs using multiple approaches.

The PITP family is composed of five members that are subdivided into two classes that exchange PI for PC (Class I, the single domain PITPα and PITPβ isoforms) or PI for phosphatidic acid (PA) (Class II, the single domain PITP cytoplasmic 1 (PITPNC1) and the multi-domain PITP membrane-associated 1 and 2 (PITPNM1 and PITPNM2) isoforms with C-terminal extensions)^25^ (Fig. 1a). The PITP-p53 interactions were first examined by immunoprecipitation (IP). Class I, but not class II PITPs, bound mutant p53 (p53^mt^), and these interactions were increased by genotoxic stress (Fig. 1b-d,f,g). Of note, the PITPα antibody had partial cross-reactivity with PITPβ but not beyond the class I PITPs (Extended Data Fig. 1a). These IP interactions were recapitulated by *in vitro* binding studies with recombinant PITPα/β and p53 proteins demonstrating a high-affinity, saturable binding of class I PITPs to p53 (Extended Data Fig. 1b-e) Moreover, the small heat shock proteins (sHSPs) Hsp27 and αB-crystallin, which stabilize the p53-PIP_n_ complex^18^, were also detected as stress-regulated components of the PITPα/β-p53 complex (Fig. 1b,e,f,h,i).

**Figure 1.**
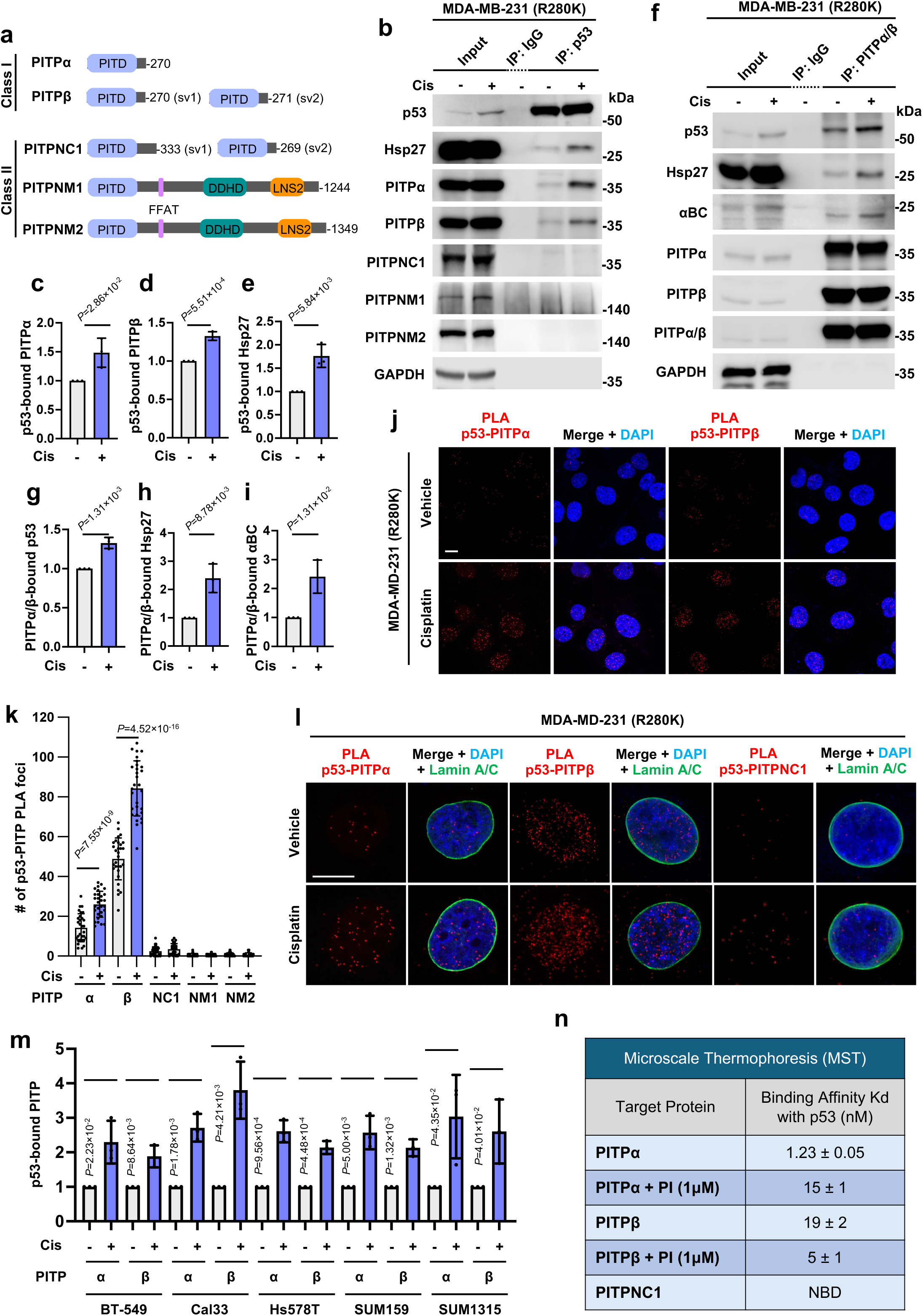
p53 interacts with the PITPs in the nucleus upon stress. **a**, Domains of PITP class I and II protein family members. PITD, PI transfer domain; FFAT, two phenylalanines in an acidic tract; DDHD, Asp-Asp-His-Asp motif; LNS2, Lipin/Nde1/Smp2 domain. **b-e**, Co-IP of PITPα/PITPβ/Hsp27 with p53 from MDA-MB-231 cells treated with vehicle or 30 µM cisplatin for 24 h. PITPα IP-ed by p53 (**c**), PITPβ IP-ed by p53 (**d**), and Hsp27 IP-ed by p53 (**e**), as well as PITPNC1/PITPNM1/PITPNM2 were analyzed by WB. *p* value denotes two-sided paired t-test. **f-i**, Co-IP of p53/Hsp27/αB-crystallin (αBC) with the shared epitope of PITPα/β from MDA-MB-231 cells treated with vehicle or 30 µM cisplatin for 24 h. p53 IP-ed by PITPα/β (**g**), Hsp27 IP-ed by PITPα/β (**h**), and αBC IP-ed by PITPα/β (**i**) were analyzed by WB. *p* value denotes two-sided paired t-test. **j-k,** PLA of p53-PITPα/PITPβ/PITPNC1/PITPNM1/PITPNM2 in MDA-MB-231 cells treated with vehicle or 30 µM cisplatin for 24 h. The nuclear p53-PITP foci were quantified (**k**). n=3, 10 cells from each independent experiment. *p* value denotes two-sided paired t-test. See expanded images in Extended Data Fig. 2a. **l,** PLA of p53-PITPα/PITPβ/PITPNC1/PITPNM1/PITPNM2 overlaid with the nuclear envelope marker Lamin A/C in MDA-MB-231 cells treated with vehicle or 30 µM cisplatin for 24 h. The nuclei were counterstained by DAPI. See expanded images in Extended Data Fig. 2e. **m,** Co-IP of PITPα/PITPβ with p53 in BT549, Cal33, HS578T, SUM159, SUM1315 cells treated with vehicle or 30 µM cisplatin for 24 h. PITPα/PITPβ IPed by p53 (**n**) and PITPNC1/PITPNM1/PITPNM2 were analyzed by WB. See expanded images in Extended Data Fig. 3j-n. **n,** The interaction of recombinant fluorescently labelled p53 with other components was quantitated by MST assay. A constant concentration of fluorescently labelled p53 (5 nM) was incubated with increasing concentrations of non-labelled ligand with or without the addition of 1 μM PI and analyzed using a Monolith NT.115 pico, and the binding affinity was autogenerated by MO. Control v.1.6 software. Data are presented as the mean ± SD. For all panels, n=3 independent experiments.

Proximity ligation assay (PLA) is an established approach to detect protein-protein interactions and posttranslational modifications (PTMs) *in situ*^30, 31^. Class I PITPs (PITPα and PITPβ), but not class II PITPs (PITPNCI, PITPNM1 and PITPNM2), associated with p53^mt^ and wild-type p53 (p53^wt^) in the nucleus as indicated by PLA foci, and their association was enhanced by cisplatin (Fig. 1j,k and Extended Data Fig. 2a-c). Interestingly, PITPα/β-p53 PLA complexes were present in the nucleoplasm in regions distinct from the nuclear envelope (Fig. 1l, Extended Data Fig. 2d-f, and Supplementary Video 1-2). Despite their previously reported confinement to the cytosol^25, 26^, PITPα/β were observed in the nucleus under basal conditions and chemotharputic stress resulted in their nuclear accumulation, supporting the PLA data (Extended Data Fig. 3a). Notably, the association of PITPα/β with p53^wt^ was minimal under basal conditions and dramatically increased by stress, while PITPα/β associated constitutively with p53^mt^ with a modest increase in response to stress. Regardless of p53 mutational status, stress-responsive PITPα/β-p53 complexes were detected in all cell lines examined (Fig. 1m and Extended Data Fig. 3b-n). Moreover, p53 directly bound class I PITPs, but not PITPNC1, with low nM K_d_ as determined by microscale thermophoresis (MST) (Fig. 1n), underscoring the specificity of these interactions.

### PITPα/β are required for p53-PIP_n_ complex generation

As PITPα/β bind to p53 in the nucleus, we examined the potential role of PITPα/β in regulating the PIP_n_ associations with p53, which stabilize p53^18^. The p53-PIP_n_ complexes^17, 18^ were first analyzed by PLA. Single or combined knock down (KD) of PITPα/β, but not PITPNC1, reduced or largely eliminated, respectively, p53^mt^-PI4,5P_2_ and p53^mt^-PI3,4,5P_3_ complexes detected by PLA; combined KD of PITPα/β disrupted p53^mt^-PIP_n_ complexes to a similar degree as p53 KD (Fig. 2a-c and Extended Data Fig. 4a,b). Similar results were observed in p53^wt^ cells: individual or combined PITPα/β KD inhibited p53^wt^-PI4,5P_2_ and p53^wt^-PI3,4,5P_3_ complexes (Fig. 2d-f).

**Figure 2.**
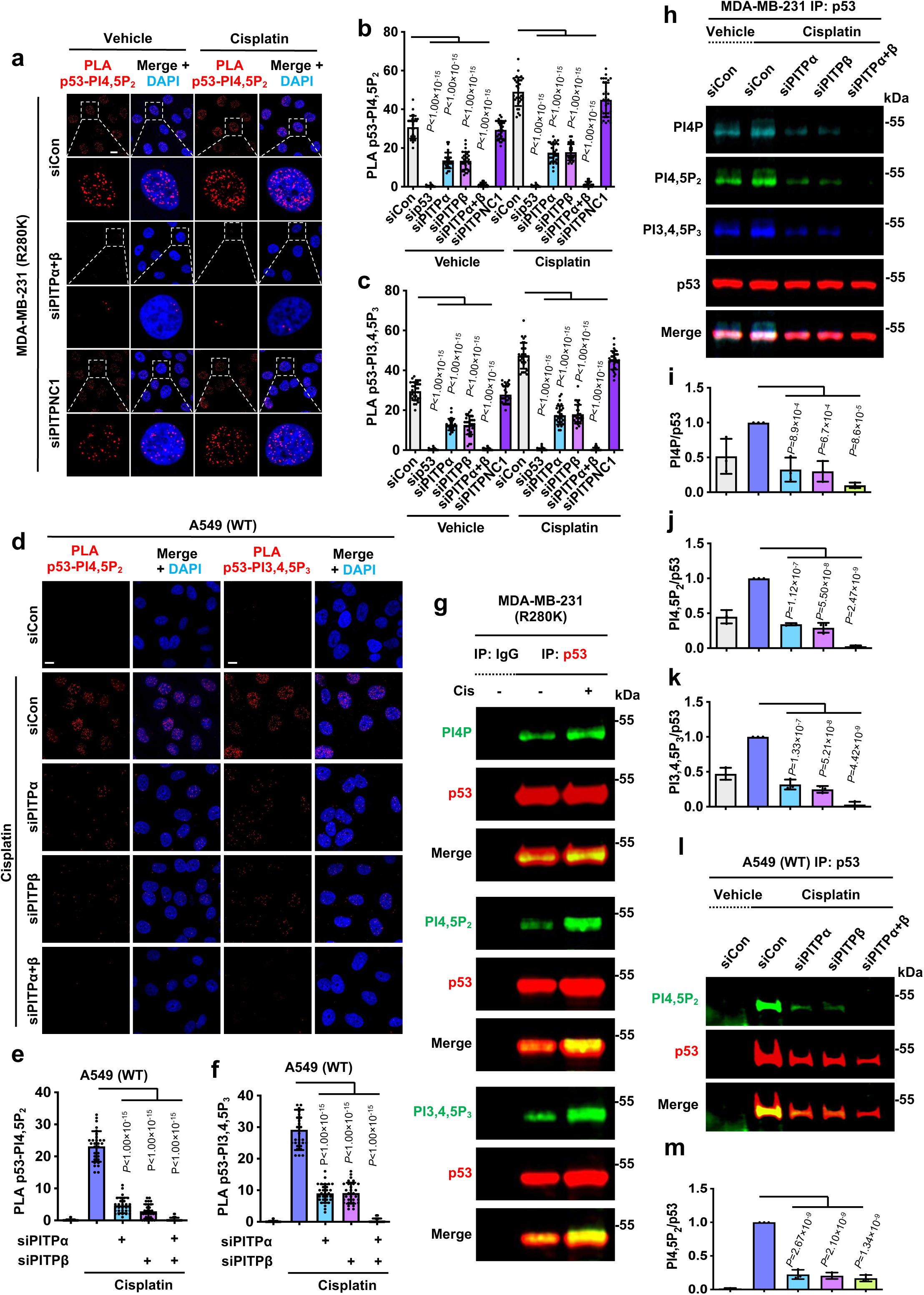
PITPα/β regulate p53-PIP_n_ complexes. **a-c,** MDA-MB-231 cells were transfected with control siRNAs or siRNAs against p53, PITPα, PITPβ, PITPNC1, or both PITPα and PITPβ. After 24 h, cells were treated with 30 µM cisplatin or vehicle for 24 h before being processed for PLA to detect p53-PI4,5P_2_/-PI3,4,5P_3_ complexes (**b,c**). n=3, 10 cells from each independent experiment. *p* value denotes ANOVA with Bonferroni’s multiple comparisons test. See KD validation in Extended Data Fig. 4d and expanded images in Extended Data Fig. 4a,b. **d-f,** A549 cells were transfected with control siRNAs or siRNAs against PITPα, PITPβ, or both PITPα and PITPβ. After 24 h, cells were treated with 30 µM cisplatin or vehicle for 24 h before being processed for PLA to detect p53-PI4,5P_2_/PI3,4,5P_3_ complexes (**e,f**). n=3, 10 cells from each independent experiment. *p* value denotes ANOVA with Bonferroni’s multiple comparisons test. See KD validation in Extended Data Fig. 4f. **g,** MDA-MB-231 cells were treated with 30 µM cisplatin or vehicle for 24 h before being processed for IP against p53. p53-bound PI4P/PI4,5P_2_/PI3,4,5P_3_ were examined by fluorescent WB. See quantification in Extended Data Fig. 4c. **h-k**, MDA-MB-231 cells were transfected with control siRNAs or siRNAs against PITPα, PITPβ, or both PITPα and PITPβ. After 24 h, cells were treated with 30 µM cisplatin or vehicle for 24 h before being processed for IP against p53. p53-bound PI4P/PI4,5P_2_/PI3,4,5P_3_ were examined by fluorescent WB and quantified by ImageJ (**i-k**). *p* value denotes ANOVA with Bonferroni’s multiple comparisons test. See KD validation in Extended Data Fig. 4d. **l-m**, A549 cells were transfected with control siRNAs or siRNAs against PITPα, PITPβ, or both PITPα and PITPβ. After 24 h, cells were treated with 30 µM cisplatin or vehicle for 24 h before being processed for IP against p53. The PI4,5P_2_ level on IPed p53 was quantified by ImageJ (**m**). *p* value denotes ANOVA with Bonferroni’s multiple comparisons test. See KD validation in Extended Data Fig. 4f. For all panels, n=3 independent experiments and data are represented as mean ± SD. Scale bar, 5 µm.

We next identified and quantified the cisplatin-induced PIP_n_ isoforms linked to p53 by p53-IP followed by simultaneous Western blotting (WB) with antibodies (Abs) against specific PIP_n_ isomers detected with fluorescently labeled secondary Abs (IP fluorescent-WB) (Fig. 2g and Extended Data Fig. 4c). KD of PITPα or PITPβ markedly reduced the PI4P, PI4,5P_2_ and PI3,4,5P_3_ that co-IPed with p53^mt^ in response to genotoxic stress, while the combined KD virtually eliminated all PIP_n_s associated with p53^mt^ (Fig. 2h-k). Similar observations were made in cells with p53^wt^: single or combined PITPα/β KD also inhibited cisplatin-stimulated p53^wt^-PI4,5P_2_ complex formation (Fig. 2l,m). Consistent with the reported role of PIP_n_s in stabilizing p53^18^, KD of PITPα or PITPβ, but not PITPNC1, modestly reduced p53^mt^ levels in cells treated with cisplatin, while the combined KD of PITPα and PITPβ robustly suppressed p53^mt^ levels under basal and stress conditions (Extended Data Fig. 4d). Reduced p53 levels after combined PITPα/β KD were not rescued by the MDM2 inhibitor Nutlin 3a^32^, suggesting a MDM2-independent mechanism (Extended Data Fig. 4e). Moreover, the robust induction of p53^wt^ by cisplatin was inhibited by individual PITPα/β KD and abrogated by combined PITPα/β KD (Extended Data Fig. 4f). The PIP_n_ antibody approach was confirmed using radiolabeled [^3^H]-inositol which is specifically incorporated into PIP_n_s and inositol phosphates^33^. [^3^H]-inositol was added to the culture media and p53 was IPed from these cells, resolved via WB, and excised from the gel, resulting in a high degree of consistency between antibody reactivity and [^3^H] counts (Fig. 3a-b and Extended Data Fig. 4g). These findings indicate that class I PITPα and PITPβ play a key role in the stress-responsive coupling of PIP_n_s to p53^wt^ and p53^mt^ proteins. The p53 associated PIP_n_, identifiable by monoclonal antibody and [^3^H]-inositol lableing marks a unique and stable protein-lipid complex that persists through denaturation and SDS-PAGE analysis.

**Figure 3.**
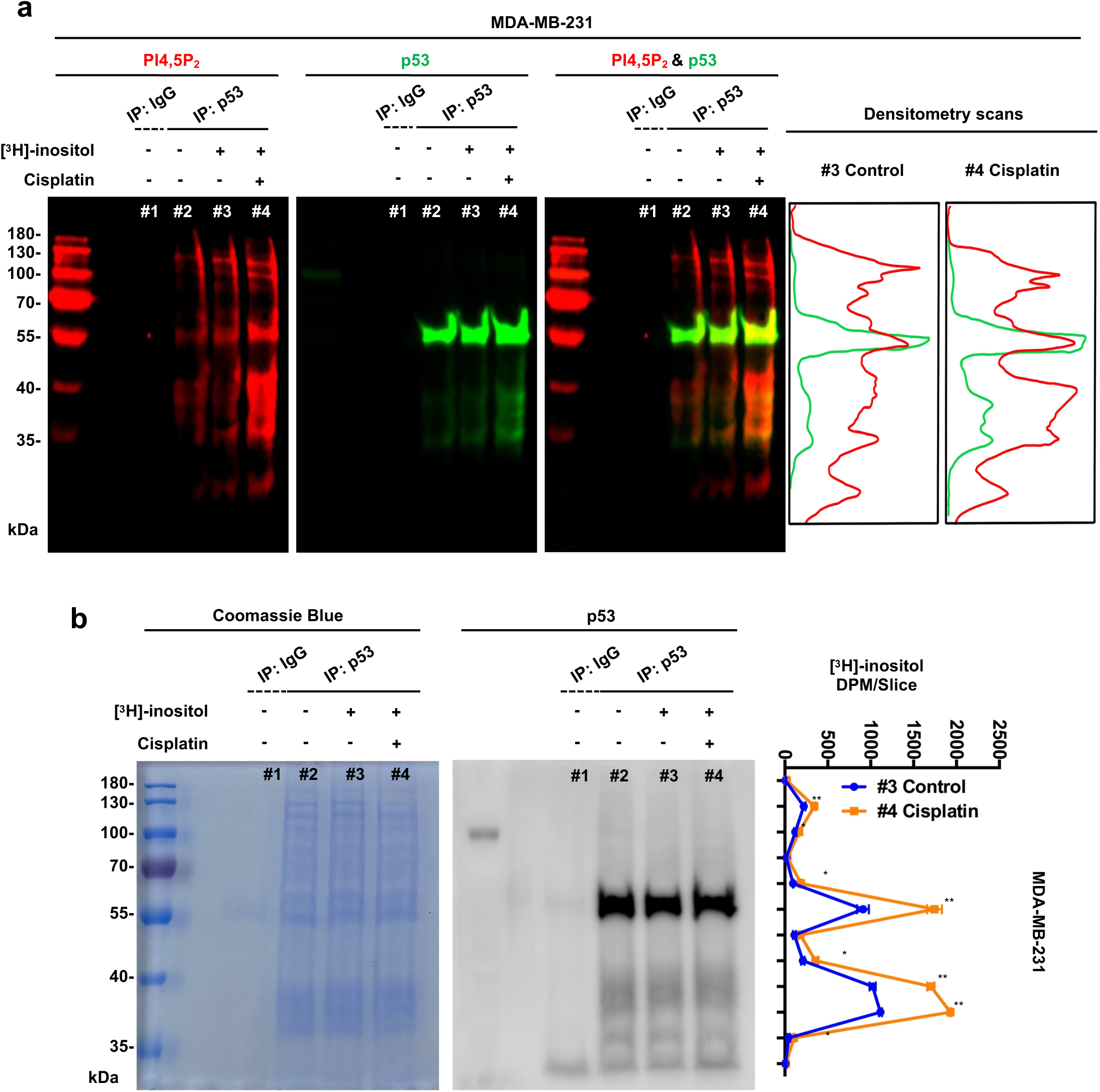
p53-PIP_n_ complexes are validated by [^3^H]-inositol lableing. **a-b,** MDA-MB-231 cells were cultured from low confluency in media containing [^3^H]*myo*-inositol or unlabeled *myo*-inositol. After 72 h of growth to confluency, cells were treated with 30 µM cisplatin or vehicle for 24 h before being processed for IP against p53. p53 and PI4,5P_2_ were confirmed by fluorescent WB (**a**) before samples were resolved by SDS-PAGE and the gel lane was excised and sectioned. Gel sections were then dissolved and analyzed by liquid scintillation counting (LSC) (**b**). n=3 independent experiments, error bar denotes SD, t-test, p<0.01*, p<0.001**. For all graphs, data are presented as the mean ± SD.

### PI4KIIα interacts with p53 and cooperates with PITPα/β to synthesize p53-PI4P

As PITPα/β were reported to cooperate with PI 4-kinases (PI4Ks) to synthesize PI4P^25, 34, 35^, we postulated that PI4Ks act in concert with PITPα/β to generate p53-PI4P. To explore this, each of the four human PI4Ks^36^ were individually KDed and PLA was used to quantify changes in the p53-PI4P complexes. KD of PI4KIIα specifically suppressed basal and cisplatin-stimulated p53-PI4P levels, while combined KD of PI4KIIα and PI4KIIIα reduced p53-PI4P levels to the same degree as PI4KIIα KD alone (Fig. 4a,b and Extended Data Fig. 5a). p53-PI4P is the substrate for PIPKIα to synthesize p53-PI4,5P_2_, which stabilizes p53^18^. Consistent with its role in generating p53-PI4P, PI4KIIα KD, but not KD of other PI4Ks, inhibited cisplatin-stimulated p53 levels (Fig. 4c). Moreover, KD of PI4KIIα mirrored the effects of PIPKIα KD in suppressing p53 induction by cisplatin, and combined KD of PI4KIIα and PIPKIα was equivalent to either individual KD with regard to p53 levels (Fig. 4d). In contrast to PI4KIIα KD which reduced p53-PI4P levels, KD of PIPKIα enhanced p53-PI4P levels (Fig. 4e,f and Extended Data Fig. 5b). p53-PI4P synthesis requires the catalytic activity of PI4KIIα as the PI4KIIα inhibitors (PI-273 and NC03)^37, 38^ prevented p53-PI4P generation and subsequent formation of p53-PI4,5P_2_ (Fig. 4g and Extended Data Fig. 5c-e). To further define these p53-PIP_n_ complexes, we utilized a p53 C-terminal domain (CTD) polybasic domain mutant (p53^6Q^), which disrupts PI4,5P_2_ binding to p53 (wild-type and oncogenic mutants)^17, 18^. In this context, the p53^6Q^ mutant formed more abundant p53^6Q^-PI4P complexes than the corresponding wild-type CTD domain p53 proteins (Extended Data Fig. 6a,b). However, the p53^6Q^-PI4P complexes were not phosphorylated further to p53^6Q^-PI3,4,5P_3_ complexes (Extended Data Fig. 6c,d), likely due to disrupted binding of PIPKIα and/or IPMK, which interact with the p53 C-terminus^17, 18^. Conventional PIP_n_ binding to p53 relies on a weak electrostatic interaction between a polybasic and positively charged surface of p53 and the acidic and negatively charged headgroup of the PIP_n_. This manner of interaction often encompasses the PIP_n_ headgroup and is ablated by harsh denaturing conditions. The p53^6Q^ mutant loses key basic amino acids and disrupts conventional PIP_n_ binding^18^. However, the stable PI4P association with p53 is enhanced by the p53^6Q^ mutation as determined by PLA, persists through denaturation as demonstrated by WB, and retains headgroup accessibility and signaling competency as previously shown^17^.

**Figure 4.**
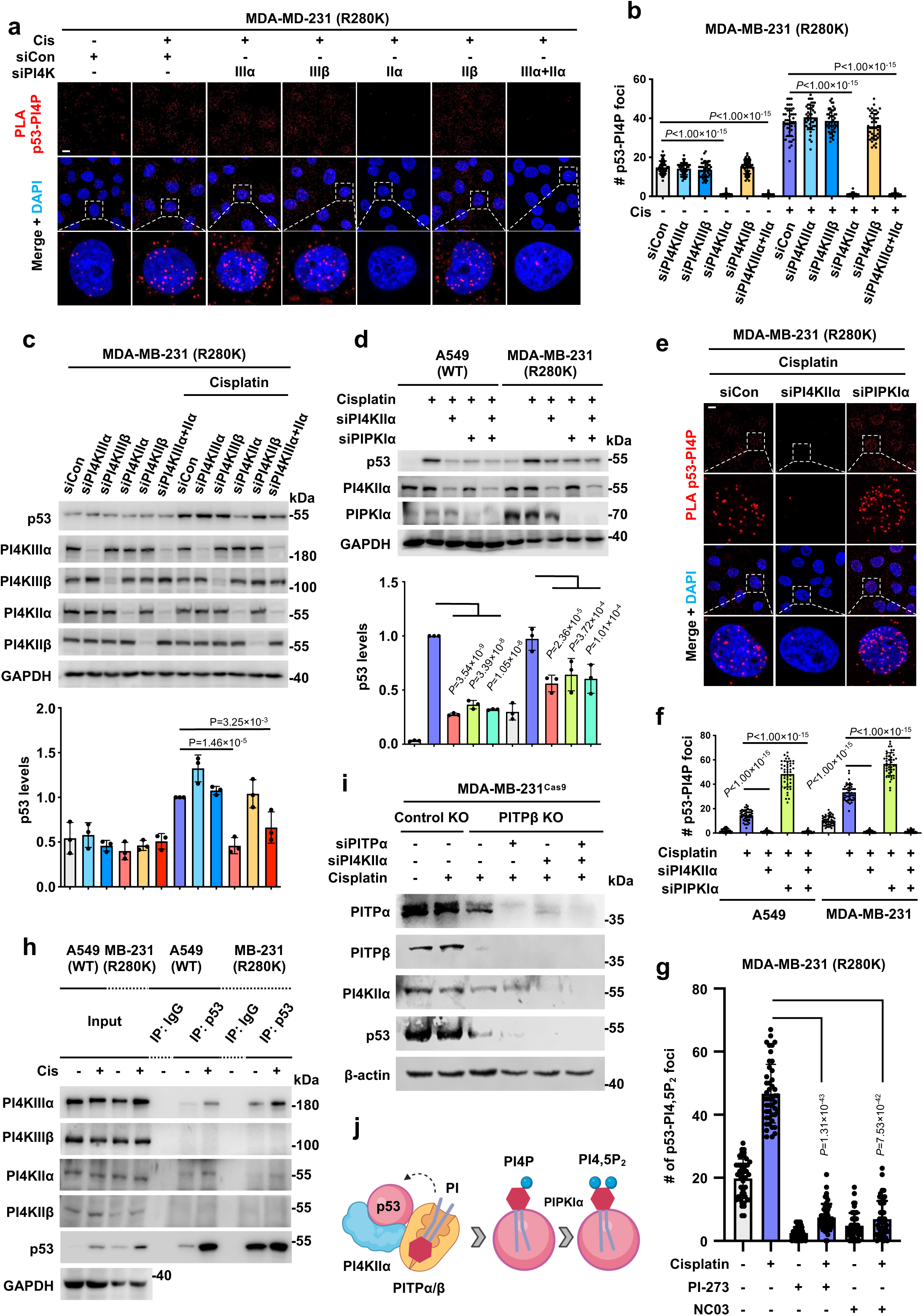
PI4KIIα interacts with p53 to generate p53-PI4P complexes. **a-b,** MDA-MB-231 cells were transfected with control siRNAs or siRNAs against PI4KIIIα, PI4KIIIβ, PI4KIIα, PI4KIIβ or both PI4KIIα and PI4KIIIα. After 24 h, cells were treated with 30 µM cisplatin or vehicle for 24 h before being processed for PLA to detect p53-PI4P complexes (**b**). n=3, 15 cells from each independent experiment. *p* value denotes ANOVA with Bonferroni’s multiple comparisons test. See KD confirmation in Fig. 4c and expanded images in Extended Data Fig. 4a. **c**, MDA-MB-231 cells were transfected with control siRNAs or siRNAs against PI4KIIIα, PI4KIIIβ, PI4KIIα, PI4KIIβ or both PI4KIIα and PI4KIIIα. After 24 h, cells were treated with 30 µM cisplatin or vehicle for 24 h before being processed for WB. p53 levels were quantified by ImageJ. *p* value denotes ANOVA with Bonferroni’s multiple comparisons test. **d,** A549 and MDA-MB-231 cells were transfected with control siRNAs or siRNAs against PI4KIIIα, PIPKIα or both PI4KIIα and PIPKIα. After 24 h, cells were treated with 30 µM cisplatin or vehicle for 24 h before being processed for WB. p53 levels were quantified by ImageJ. *p* value denotes ANOVA with Bonferroni’s multiple comparisons test. **e,f,** MDA-MB-231 and A549 cells were transfected with control siRNAs or siRNAs against PI4KIIIα, PIPKIα or both PI4KIIα and PIPKIα. After 24 h, cells were treated with 30 µM cisplatin or vehicle for 24 h before being processed for PLA to detect p53-PI4P complexes. n=3, 15 cells from each independent experiment. *p* value denotes ANOVA with Bonferroni’s multiple comparisons test. See KD confirmation in Fig. 4d and expanded images in Extended Data Fig. 4b. **g**, MDA-MB-231 cells were treated with vehicle or cisplatin in combination with DMSO as a control, PI-273, or NC03 for 24 h before being processed for PLA to detect p53-PI4,5P_2_ complexes and analyzed using ImageJ. *p* value denotes two-sided paired t-test. n=3, 15 cells from each independent experiment. See expanded images in Extended Data Fig. 5e. **h**, MDA-MB-231 and A549 cells were treated with 30 µM cisplatin or vehicle for 24 h before being processed for IP against p53. The PI4KIIIα, PI4KIIIβ, PI4KIIα, and PI4KIIβ levels IPed with p53 were analyzed by WB. **i**, MDA-MB-231^Cas9^ cells with PITPβ KO and control non-targeted KO were transfected with control siRNAs or siRNAs against PI4KIIα, PITPα, or both PI4KIIα and PITPα. After 24 h, cells were treated with 30 µM cisplatin or vehicle for 24 h before being processed for WB. **j,** Model showing PITPα/β cooperation with PI4KIIα in generating p53-PI4P complexes that participate in further signaling. For all panels, n=3 independent experiments and data are represented as mean ± SD. Scale bar, 5 µm.

In addition, PI4KIIα co-IPed with p53^mt^ and p53^wt^, and these interactions were enhanced by genotoxic stress (Fig. 4h). PI4KIIIα also co-IPed with p53^mt^ and p53^wt^, (Fig. 4h) but PI4KIIIα KD did not affect p53 or p53-PI4P levels or enhance the effects of PI4KIIα KD (Fig. 4a-c). Recombinant PI4KIIα bound to p53, PITPα, and PITPβ with saturatable kinetics and low nM affinity (Extended Data Fig. 7a-d), consistent with a specific and direct interaction. Interestingly, the PI4KIIα-PITPβ interaction was enhanced by the addition of p53, suggesting a ternary PI4KIIα:PITPβ:p53 protein complex that contributes to p53 stability (Fig. 4i and Extended Data Fig. 7e-g). These findings indicate that PI4KIIα binds to p53 with high affinity and, together with PITPα/β, specifically synthesizes p53-PI4P and contributes to p53 stabilization by generating the precursor for p53-PI4,5P_2_ (Fig. 4j).

### PITPα/β regulate p53-PIP_n_ signalosome formation and nuclear Akt activation to inhibit stress-induced apoptosis

The p53-PIP_n_ signalosome is regulated by cellular stress, leading to the formation of p53-PI3,4,5P_3_ complexes that recruit and activate Akt in the nucleus and protect cells from stress-induced apoptosis^17^. The identification of class I PITPs as upstream regulators of p53-PIP_n_ complex assembly implicates these lipid transfer proteins in nuclear Akt activation. Consistent with this concept, individual KD of PITPα or PITPβ, but not PITPNC1, suppressed stress-induced nuclear Akt activation quantified by nuclear pAkt^S473^ IF, an established approach^17^, while combined PITPα/β KD resulted in a more robust reduction of pAkt^S473^ (Fig. 5a,b). These results were confirmed in MDA-MB-231 cells by WB (Extended Data Fig. 8a). Similar results were obtained with multiple human cancer cell lines expressing different p53^mt^ proteins (Fig. 5c-e). KD of p53^mt^, PITPα or PITPβ, but not PITPNC1, moderately reduced cell viability under basal conditions and sensitized MDA-MB-231 cells to cisplatin treatment, while combined KD of PITPα/β further decreased cell viability (Fig. 5f,g). Consistent with these results, knockout (KO) of PITPβ or KD of PITPα had modest effects on cisplatin-induced cell death, while combined PITPβ KO and PITPα KD enhanced the cytotoxicity of cisplatin (Fig. 5h and Extended Data Fig. 8b). Individual or combined PITPα/β KD had similar effects on cisplatin sensitivity in four additional breast cancer cell lines expressing different p53^mt^ or p53^wt^ proteins (Extended Data Fig. 8c-j). The cytotoxicity of p53^mt^ KD and individual or combined PITPα/β KD was associated with a corresponding increase in caspase 3 activity (Fig. 5i and Extended Data Fig. 8k-m), indicating that PITPα/β KD triggers apoptosis and sensitizes cells to genotoxic stress. These findings underscore the critical role of PITPα/β in regulating nuclear Akt activation, which confers resistance to genotoxic stress-induced apoptosis.

**Figure 5.**
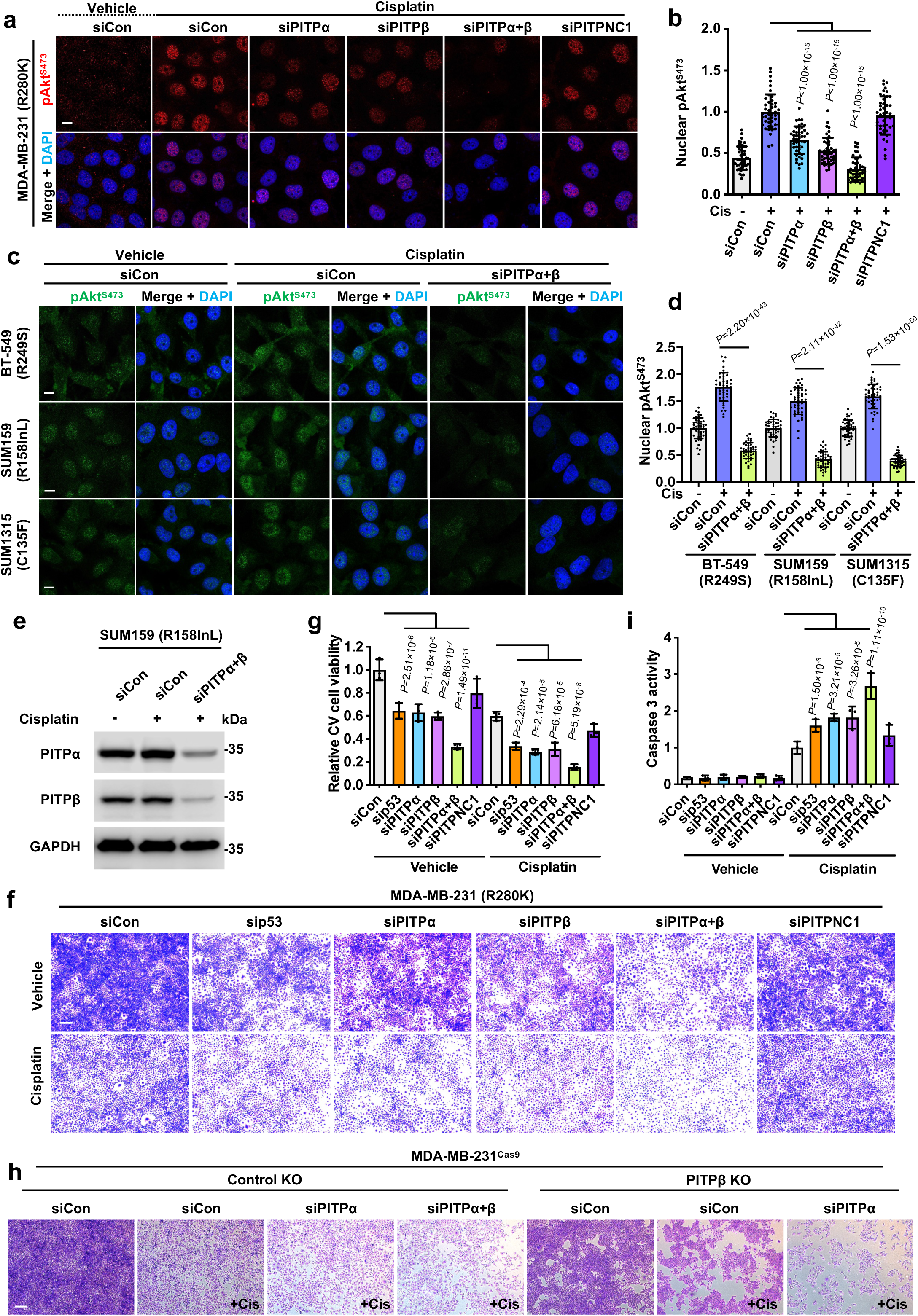
PITPα/β regulate p53-dependent nuclear Akt activation and cell survival. **a-b**, MDA-MB-231 cells were transfected with control siRNAs or siRNAs against PITPα, PITPβ, PITPNC1, or both PITPα and PITPβ. After 24h, cells were treated with 30 µM cisplatin or vehicle for 24 h before being processed for IF staining against pAkt^S473^. The nuclear pAkt^S473^ levels were quantified by ImageJ (**b**). n=3, 15 cells from each independent experiment. *p* value denotes ANOVA with Bonferroni’s multiple comparisons test. Scale bar, 5 µm. See KD validation in Extended Data Fig. 4d. **c-e**, BT-549, SUM159, and SUM1315 cells with the indicated p53 status were transfected with control siRNAs or siRNAs against both PITPα and PITPβ. After 24 h, cells were treated with 30 µM cisplatin or vehicle for 24 h before being processed for IF staining against pAkt^S473^. The nuclei were counterstained by DAPI. The nuclear pAkt^S473^ levels were quantified by ImageJ (**d**). n=3, 15 cells from each independent experiment. *p* value denotes two-sided paired t-test. Scale bar, 5 µm. The double KD of PITPα and PITPβ in SUM159 cells was validated by WB (**e**). See KD validation for BT-549 and SUM1315 cells in Extended Data Fig. 8h, j. **f-g**, MDA-MB-231 cells were transfected with control siRNAs or siRNAs against p53, PITPα, PITPβ, PITPNC1, or both PITPα and PITPβ. After 24 h, cells were treated with 30 µM cisplatin or vehicle for 24 h before being processed for Crystal Violet viability assay. The cells were imaged by an EVOS M5000 microscope (**f**) and quantified based on the extracted dye using a plate reader (**g**). *p* value denotes ANOVA with Bonferroni’s multiple comparisons test. Scale bar, 300 µm. **h,** MDA-MB-231^Cas9^ cells with PITPβ KO and control non-targeted KO were transfected with control siRNAs or siRNAs against PITPα, PITPβ, or both PITPα and PITPβ. After 24 h, cells were treated with 30 µM cisplatin or vehicle for 24 h before being processed for Crystal Violet viability assay. The cells were imaged by an EVOS M5000 microscope. Scale bar, 300 µm. See quantification in Extended Data Fig. 8b. **i**, MDA-MB-231 cells were transfected with control siRNAs or siRNAs against p53, PITPα, PITPβ, PITPNC1, or both PITPα and PITPβ. After 24h, cells were treated with 30 µM cisplatin or vehicle for 24 h before being processed for a caspase 3 activity assay. *p* value denotes ANOVA with Bonferroni’s multiple comparisons test. For all panels, n=3 independent experiments and data are presented as the mean ± SD.

### PITPα/β-mediated p53-PIP_n_ signaling requires PI transfer activity

As PITPα/β exchange PI for PC, we investigated whether the PI transfer activity of these PITPs was required to regulate p53 levels and nuclear Akt activation. For this, we utilized the PITPα/β^T59D^ mutant that specifically disrupts PI binding^39–41^. Ectopically expressed HA-tagged PITPα/β^wt^ interacted with p53^wt^ and mutant p53^R175H^, while the corresponding PITPα/β^T59D^ mutants exhibited impaired interaction with p53 (Fig. 6a and Extended Data Fig. 9a-c). Moreover, the reduction in the stress-induced p53 protein and nuclear pAkt^S473^ levels by combined PITPα and PITPβ KD using distinct siRNAs targeting their 3’-UTR was rescued by expression of PITPα^wt^, but not mutant PITPα^T59D^ (Fig. 6b-d). Notably, nuclear pAkt^S473^ levels correlated with the content of re-expressed PITPα^wt^, but not mutant PITPα^T59D^ (Fig. 6e,f), indicating that these phenotypes are dose-dependent.

**Figure 6.**
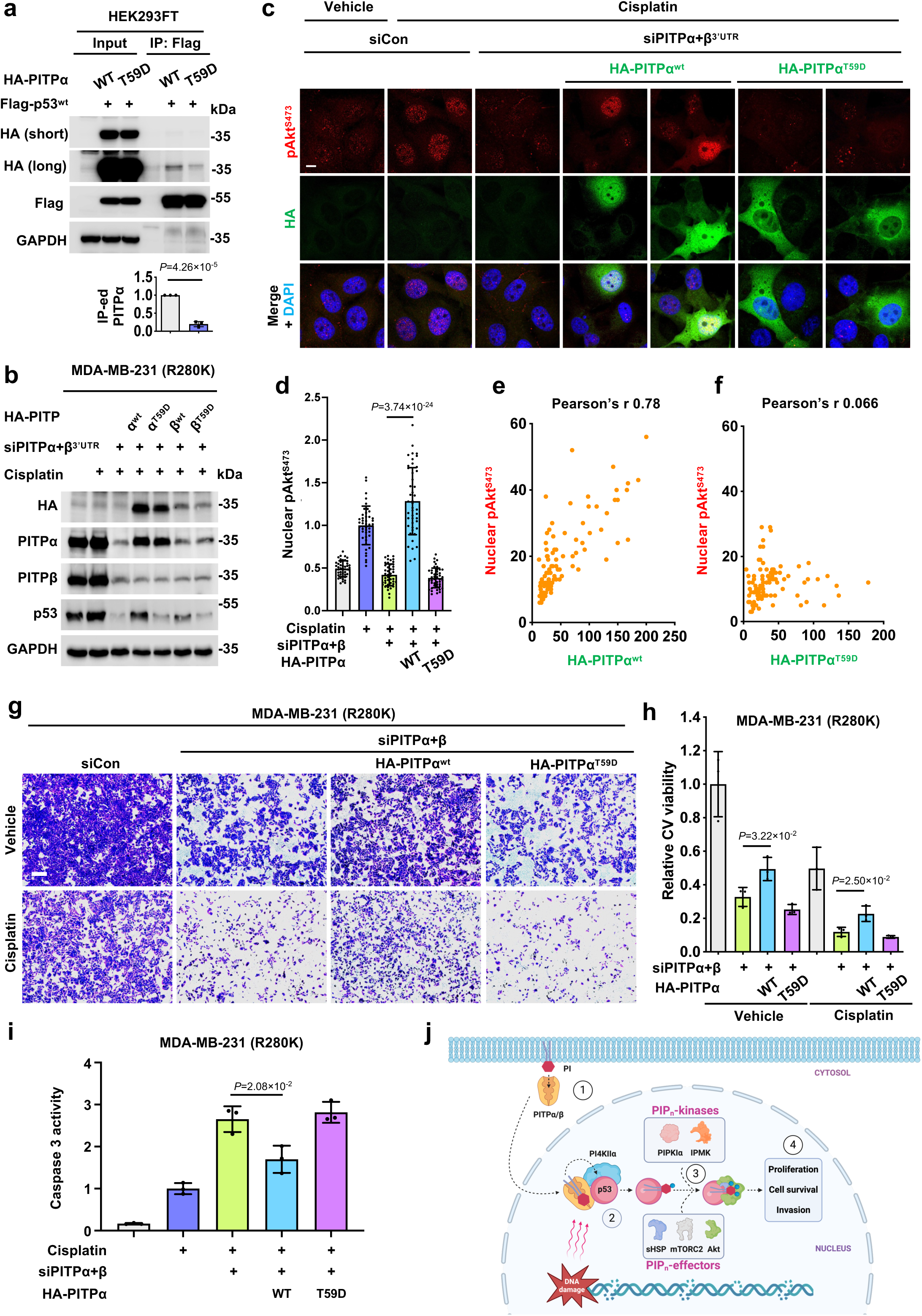
PITPα/β initiate the p53-PIP_n_ signalosome by a PI binding-dependent mechanism. **a**, HEK293FT cells were co-transfected with either HA-tagged wild-type PITPα or PI binding-defective mutant T59D PITPα together with Flag-tagged p53^wt^. After 48 h, cells were processed for IP against Flag-tag and analyzed by WB. *p* value denotes two-sided paired t-test. **b-d**, MDA-MB-231 cells were transfected with control siRNAs or siRNAs against the 3’UTR of both PITPα and PITPβ for 24 h. Then, the cells were transfected with either HA-tagged wild-type PITPα/β or PI binding-defective mutant T59D PITPα/β. After 24 h, cells were treated with 30 µM cisplatin or vehicle for 24 h before being processed for WB (**b**) and IF staining against HA-tag with pAkt^S473^ (**c**). The nuclear pAkt^S473^ levels in HA-positive cells were quantified by ImageJ (**d**). n=3, 15 cells from each independent experiment. *p* value denotes two-sided paired t-test. Scale bar, 5 µm. **e-f**, Pearson’s correlation coefficient (Pearson’s r) quantified the correlations between nuclear pAkt^S473^ and HA-PITPα^wt/T59D^ levels. n=100 cells pooled from 3 independent experiments. **g-h**, MDA-MB-231 cells were transfected with control siRNAs or siRNAs against the 3’UTR of both PITPα and PITPβ for 24 h. The cells were then transfected with either HA-tagged wild-type PITPα or PI binding-defective mutant T59D PITPα. After 24 h, cells were treated with 30 µM cisplatin or vehicle for 24 h before being processed for Crystal Violet viability assay. The cells were imaged by an EVOS M5000 microscope (**g**) and quantified based on the extracted dye using a plate reader (**h**). *p* value denotes two-sided paired t-test. Scale bar, 300 µm. **i**, MDA-MB-231 cells were transfected with control siRNAs or siRNAs against the 3’UTR of both PITPα and PITPβ for 24 h. The cells were then transfected with either HA-tagged wild-type PITPα or PI binding-defective mutant T59D PITPα. After 24 h, cells were treated with 30 µM cisplatin or vehicle for 24 h before being processed for a caspase 3 activity assay. *p* value denotes two-sided paired t-test. **j**, A model of the stress-induced nuclear accumulation of class I PITPs and PITP-dependent initiation of the p53-PI signalosome, which promotes cell proliferation, survival, and invasion. For all panels, n=3 independent experiments and data are presented as the mean ± SD.

The cytotoxicity and sensitization to cisplatin-induced apoptosis by combined PITPα/β KD using the 3’-UTR targeting siRNAs were partly rescued by re-expressing PITPα^wt^ but not the PITPα/β^T59D^ mutant (Fig. 6g-i). These results demonstrate that the PI transfer activity of class I PITPs regulates their interaction with p53, p53 stability, and subsequent nuclear Akt activation. Combined PITPα/β KD also inhibited EGF-stimulated cell invasion, and these effects were partly restored by re-expressing PITPα^wt^ but not the PITPα^T59D^ mutant (Extended Data Fig. 9d,e), consistent with reports that PITPs contribute to EGF signaling^42^. The data indicate that PITPα/β regulate cell growth, invasion, and protection against genotoxic stress (Fig. 6j), although elucidation of the underlying mechanisms will require additional investigation.

## Discussion

Here, we reveal an unexpected role for class I PITPα/β in generating nuclear p53-PIP_n_ complexes that initiate nuclear PIP_n_ signaling. Although the existence of nuclear PIP_n_s and their role in the cell cycle^43^, oxidative stress^16, 44–49^, and the DNA damage response^17, 50–52^ is supported by growing evidence^16^, the molecular regulators of nuclear PIP_n_s in regions devoid of membranes have been enigmatic^53^. PITPs have a well-established role in transporting PI from the endoplasmic reticulum to membranes, enabling membrane-localized PIP_n_ signaling via PIP kinases, phosphatases, and phospholipases^25, 54–58^. The findings presented here demonstrate that class I PITPs directly bind p53 with high affinity in the nucleus in response to genotoxic stress. As such, these data expand the roles of the PITPα/β lipid transport activities that were thought to act solely at cytosolic membranes and establish their mechanistic roles in generating nuclear p53-PIP_n_ complexes.

PITPα/β, but not other PITP family members, bind directly to p53^wt^ and p53^mt,^ and these interactions are stimulated by cell stress. This complex also includes the small heat shock proteins (Hsp27 and αB-crystallin), which stabilize p53^18^. PITPα/β binding to p53 is necessary for the linkage of PIP_n_s with p53 (p53-PIP_n_ signalosome). Once the p53-PIP_n_ complex is formed, it is stable, detected by PIP_n_ isomer antibodies, and acted on by multiple PIP kinases and phosphatases, indicating that the myo-inositol head group is accessible, a characteristic that distinguishes these complexes from conventional protein-PIP_n_ binding^17, 18, 31^. The p53-PIP_n_ signalosome culminates in the formation of p53-PI3,4,5P_3_ complexes that, in turn, recruit and activate Akt in the nucleus, conferring resistance to stress-induced apoptosis^17^. However, PITPs also contribute to the canonical membrane-localized PI3K/Akt pathway^59^, so the observed differences in cellular phenotypes by PITPα/β KD likely reflect some contribution of both the canonical and the nuclear PI3K/Akt pathways. These PITPα/β activities require PI binding, underscoring the critical functional role of lipid transport as a mechanism to establish nuclear PIP_n_s.

The ability of wild-type PITPα, but not the PI binding-defective mutant, to rescue the phenotype of combined PITPα/β KD suggests a functional redundancy between the class I PITPs. This is consistent with previous reports that either PITPα or PITPβ is sufficient to rescue the phenotype of double PITPα/β knockout in a PI-binding dependent manner^39, 60^. PITPα/β also binds to PI4KIIα, and the addition of p53 protein increases the affinity between PITPs and PI4KIIα, which supports formation of a ternary complex. As both PITPα/β and PI4KIIα are required to generate p53-PI4P, these data point to a functional collaboration between PITPα/β and PI4KIIα in initiating nuclear p53-PIP_n_ signaling that is reminiscent of their roles in generating PI4P in the membrane-localized pathway^34, 35^.

Remarkably, PITPα/β and PI4KIIα regulate the association of PIP_n_s to p53 in a manner that is inconsistent with traditional protein-PIP_n_ interactions. Conventional protein-PIP_n_ interactions are electrostatic in nature and are disrupted by denaturation and SDS-PAGE, yet p53-PIP_n_ complexes remain intact following these interventions^17, 18^. Previous studies have demonstrated that the p53^6Q^ CTD mutant has diminished binding to PIP_n_s^17, 18^, but the p53^6Q^-PI4P complex is enhanced, further distinguishing these complexes from conventional p53-PIP_n_ binding. Perhaps most significantly, the p53-linked PIP_n_s maintain signaling competency as they are modified by kinases and phosphatases and remain detectable via antibody recognition of their *myo*-inositol headgroup or [^3^H]-inositol radiolabeling. In contrast to p53-PIP_n_ complexes, most protein-PIP_n_ interactions involve direct binding of the headgroup to protein domains precluding the lipid from kinase or phosphatase modification^61, 62^, underscoring this unique characteristic of p53-PIP_n_ complexes. The data presented here indicate that PIP_n_ second messengers, beginning with PI4P, are linked to p53 by the combined activities of PITPα/β and PI4KIIα. The resulting p53-PI4P is acted on by PIP kinases and phosphatases such that the p53-PI3,4,5P_3_ functions as a nuclear messenger to recruit PDK1, mTORC2, and Akt. Notably, each of these proteins is known to interact with the PIP_n_ head group. In the case of mutant p53 proteins, p53 activation may contribute to its oncogenic gain-of-function activity by activating nuclear Akt and suppressing apoptosis, whereas this prosurvival activity may counteract the proapoptotic function of wild-type p53 and represent a novel mechanism for wild-type p53 to confer stress-resistance consistent with prior reports of its ability to regulate metabolism, oxidative stress and DNA damage^63^. These findings suggest a “third messenger” pathway^17^ with p53 as the founding member whereby PIP_n_ second messengers are linked to cellular proteins in membrane-free regions and modified by PIP kinases/phosphatases to regulate their function (Fig. 6j). Given the critical role of class I PITPs in initiating nuclear phosphoinositide signaling, which attenuates cellular stress and apoptosis, these findings also point to these PITPs and PI4KIIα as potential therapeutic targets in a variety of diseases including cancer.

## ACKNOWLEDGMENTS

We thank Dr. Adrea Galmozzi for discussions and comments and Lance Rodenkirch for technical support. N.D.C. is supported by a National Institutes of Health T32 Training Grant 5T32ES007015-43 to the Molecular and Environmental Toxicology graduate training program at UW-Madison. M.C. is supported by grants 32400577, 2023A1515110237, JCYJ20240813094605008, and D2301007. This work was supported in part by a National Institutes of Health grant R35GM134955 (R.A.A.), R01CA286492 (R.A.A. & V.L.C.), Department of Defense Breast Cancer Research Program grants W81XWH-21-1-0129 (V.L.C.), HT9425-23-1-0553 (V.L.C.), and HT9425-23-1-0554 (R.A.A.), and a grant from the Breast Cancer Research Foundation (V.L.C.).

## AUTHOR CONTRIBUTIONS

M.C., N.D.C, T.W., V.L.C., and R.A.A. designed the experiments. N.D.C., M.C., T.W., P.A., T.J.W., C.S., and D.B. performed the experiments. N.D.C., M.C., T.W., V.L.C., and R.A.A. wrote the manuscript.

## DECLARATION OF INTERESTS

Authors declare that they have no competing interests.

## Experimental Procedures

### Cell culture and constructs

A549, BT-549, Cal33, HCT116, HS578T, MDA-MB-231, SUM159, SUM1315, HEK293FT, and MCF-10A cells were purchased from ATCC. MCF10A cells were cultured in DMEM/F12 (#11330-032, Invitrogen) supplemented with 10% fetal bovine serum (#SH30910.03, Hyclone), 1% penicillin/streptomycin (#15140-122, Gibco), 20 ng/mL EGF (#CC-4107, Lonza), 0.5 mg/mL Hydrocortisone (#H4001, Sigma), 100 ng/mL Cholera toxin (#C-8052, Sigma), and 10 μg/mL Insulin (#I-1882, Sigma). The other cells were maintained in DMEM (#10-013-CV, Corning) supplemented with 10% fetal bovine serum (#SH30910.03, Hyclone) and 1% penicillin/streptomycin (#15140-122, Gibco). The Flag-tagged p53 constructs and corresponding PIP_n_-binding-defective 6Q mutants in this work have been described previously^18^. The HA-tagged wild-type PITPα/β constructs and PI-binding deficient mutants T59D^39^ were purchased from Genscript. Plasmids were transfected into mammalian cells using Lipofectamine^TM^3000, (#L3000015, Thermo Fisher Scientific) according to the manufacturer’s instructions. Typically, 2-5 µg of DNA and 6-10 µl of lipid were used for transfecting cells in 6-well plates. Cells with at least 80% transfection efficiency were used for further analysis.

### Antibodies and reagents

Monoclonal antibodies against p53 (clone DO-1, #SC-126, Santa Cruz Biotechnology), p53 (clone 7F5, #2527, Cell Signaling), pAkt^S473^ (clone 193H12, #4058, Cell Signaling), HA-tag (clone C29F4, #3724, Cell Signaling), Flag-tag (clone D6W5B, #14793, Cell Signaling), GAPDH (clone 0411, #sc-47724, Santa Cruz Biotechnology), Lamin B2 (clone D8P3U, #12255, Cell Signaling), EGFR (clone D6B6, #2085, Cell Signaling), HSP27 (clone D6W5V, #95357, Cell Signaling), and polyclonal antibodies against PITPα (#16613-1-AP, ThermoFisher), PITPβ (#ab127563, abcam), PITPNC1 (IF, #ab222078, abcam), PITPNC1 (WB, #NBP2-19842, Novus), PITPNM1 (#26983-1-AP, Proteintech), PITPNM2 (#PA5-48616, Invitrogen), pAkt^T308^ (#9275, Cell Signaling), PI4KIIα (#NBP2-44158, Novus), PI4KIIIα (#4902S, CST), PI4KIIβ (#PA5-82133, Thermo), PI4KIIIβ (#ab134756, abcam), and αB-Crystallin (#ADI-SPA-223-F, Enzo) were used in this study. For conventional immunostaining and PLA analysis of phosphoinositides, anti-PI4P (#Z-P004), PI4,5P_2_ (#Z-P045), and PI3,4,5P_3_ (#Z-P345) antibodies were purchased from Echelon Biosciences. For immunoblotting analyses, antibodies were diluted at a 1:1000 ratio except for p53 (clone DO-1, 1:5000) and GAPDH (clone 0411, 1:5000). For immunoprecipitation, antibody-conjugated agarose was purchased from Santa Cruz Biotechnology, including agarose-conjugated antibodies against p53 (#sc-126AC), PITPα/β (#sc-398050AC), HA-tag (#sc-7392AC), and Flag-tag (#sc-166355AC). For immunostaining analyses and proximity ligation assay (PLA), antibodies were diluted at a 1:100 ratio. The nuclear envelope marker (Alexa Fluor**®**488 Lamin A/C, clone 4C11, #8617, Cell Signaling, 1:200) examined subnuclear regions. For the knockdown (KD) experiments, the ON-TARGETplus siRNA SMARTpool with four siRNAs in combination against human p53 (#L-003329-00), PITPα (#L-018010-00), PITPβ (#L-006459-01), and PITPNC1 (#L-012476-02) were purchased from Dharmacon. Non-targeting siRNA (#D-001810-01, Dharmacon) was used as a control. For the KD and rescue experiments, pooled-siRNAs targeting the 3’UTR of PITPα (sense 5’-AGGCAACUUUCGAUUCCUUACUGUA-3’ and antisense 5’-UACAGUAAGGAAUCGAAAGUUGCCU-3’; sense 5’-CCGUCUCUCUCCAUUGUGUUCCGAU-3’ and antisense 5’-AUCGGAACACAAUGGAGAGAGACGG-3’) and pooled-siRNAs targeting the 3’UTR of PITPβ (sense 5’-GAGUGAACAACAAUCUGACCAGUAU-3’ and antisense 5’-AUACUGGUCAGAUUGUUGUUCACUC-3’; sense 5’-CUACAAAGCUGAUGAAGAC-3’ and antisense 5’-GUCUUCAUCAGCUUUGUAG-3’) were purchased from Thermo Fisher Scientific. The siRNAs were delivered to cells by RNAiMAX reagent (#13778150, Thermo Fisher Scientific), and KD efficiency was determined by immunoblotting. KD efficiency greater than 80% was required to observe phenotypic changes in the study. Cisplatin (#NC1706394, Fisher Scientific), etoposide (#S1225, Sellekchem), Erastin (#S7242, Selleckchem), tert-Butylhydroquinone (tBHQ, #SPCM-T1540-06, Spectrum), and hydroxyurea (HU, #1332000, Millipore Sigma) were used as cellular stressors. Nutlin 3a (#S8059, Sellekchem), PI-273 (#AOB37765, aobious), and NC03 (#AOB17420, aobious) were also used.

### Immunoprecipitation and immunoblotting

Cells were removed from the medium after the indicated treatment, washed three times with ice-cold PBS, and lysed in an ice-cold RIPA lysis buffer system (#sc-24948, Santa Cruz Biotechnology) with 1 mM Na_3_VO_4_, 5 mM NaF, and 1x protease inhibitor cocktail (#11836153001, Roche). The cell lysates were then sonicated followed by incubation at 4°C with continuous rotation for 1 h and centrifuged at maximum speed for 10 min to collect the supernatant. The protein concentration in the supernatant was measured by the Bradford protein assay (#5000201, BIO-RAD). All antibodies were diluted at a 1:1000 ratio for immunoblotting unless otherwise indicated. For immunoprecipitation, 0.5-1 mg of protein was incubated with 20 µl of antibody-conjugated agarose (Santa Cruz Biotechnology) at 4°C for 24 h. After washing three times with PBST (PBS with 0.1% Tween 20), the protein complex was eluted with SDS sample buffer and boiled at 95°C for 10 min. For immunoblotting, 5-20 µg of protein were loaded. For immunoblotting of immunoprecipitated complexes, horseradish peroxidase (HRP)-conjugated antibodies were used to avoid non-specific detection of immunoglobulin in the immunoprecipitated samples. HRP-conjugated p53 (#sc-126HRP), PITPα (#sc-13569HRP), and PITPβ (#sc-390500HRP) antibodies were purchased from Santa Cruz Biotechnology. Immunoblots were developed by Odyssey Imaging System (LI-COR Biosciences), and the intensity of protein bands was quantified using ImageJ. Statistical data analysis was performed with Microsoft Excel, using data from at least three independent experiments.

### Fluorescent IP-WB

Cells were lysed in a RIPA lysis buffer system^17^ and lysates were incubated with 20 µl anti-p53 (#sc-126 AC, Santa Cruz Biotechnology) mouse monoclonal IgG antibody-conjugated agarose at 4°C for 24 h. Normal immunoglobulin (IgG)-conjugated agarose was used as a negative control (#sc-2343, Santa Cruz Biotechnology). The protein complex was eluted with SDS sample buffer and boiled at 95°C for 10 min. For immunoblotting, 5-20 µg of protein were loaded. The protein complexes were analyzed via WB as described above. For double fluorescent IP-WB detecting p53-PI4P/PI4,5P_2_/PI3,4,5P_3_ complexes, anti-p53 rabbit monoclonal IgG antibody (clone 7F5, #2527, Cell Signaling) at 1:2000 dilution and anti-PI4P mouse monoclonal IgM antibody (#Z-P004, Echelon Biosciences), PI4,5P_2_ mouse monoclonal IgM antibody (#Z-P045, Echelon Biosciences), or PI3,4,5P_3_ mouse monoclonal IgM antibody (#Z-P345, Echelon Biosciences) at 1:1000 dilution were mixed in blocking buffer with 0.02 % Sodium Azide and incubated with the membrane at 4°C overnight. For the secondary antibody incubation, goat anti-rabbit IgG antibody conjugated with IRDye 800CW fluorophore (#926-32211, LI-COR) and goat anti-mouse IgM antibody conjugated with IRDye 680RD fluorophore (#926-68180, LI-COR) at 1:10000 dilution were mixed in blocking buffer with 0.01 % SDS and 0.1% Tween 20 and incubated with the membrane at room temperature for 2 h. The images were subsequently acquired simultaneously using the 700 and 800 nm wavelength channels on the Odyssey Fc Imaging System (LI-COR Biosciences). Statistical data analysis was performed with Microsoft Excel, using data from at least three independent experiments.

### Immunofluorescence (IF) and Confocal Microscopy

For immunofluorescence studies, cells were grown on coverslips coated with 0.2 % gelatin (#G9391, Millipore Sigma). Cells were fixed with 4% paraformaldehyde (PFA) (#sc-281692, Santa Cruz Biotechnology) for 20 min at room temperature, then washed three times with PBS. Next, the cells were permeabilized with 0.3% Triton-X100 for 30 min to permeate the nuclear envelope thoroughly and were rewashed three times with PBS. The cells were then blocked with 1% BSA in PBS for one hour at room temperature. After blocking, cells were incubated with primary antibodies overnight at 4°C. The cells were then washed three times with PBS and incubated with fluorescent-conjugated secondary antibodies (Molecular Probes) for 1 hour at room temperature. After secondary antibody incubation, the cells were washed three times. Then the coverslips were removed from the dishes, air-dried for 10 min at room temperature and mounted in Prolong^TM^ Glass Antifade Mountant with NucBlue^TM^ Stain (#P36985, Thermo Fisher Scientific). The images were taken by a Leica SP8 3xSTED Super-Resolution Microscope, a point scanning confocal, and a 3xSTED super-resolution microscope using LASX software (Leica Microsystems). All images were acquired using the 100X objective lens (N.A. 1.4 oil). The z-stack images were taken with each frame of 0.2 µm thickness. For quantification, the mean fluorescent intensity of channels in each cell was measured by LASX. GraphPad Prism Version 9.4.1 generated quantitative graphs. The images were processed using ImageJ. The correlation of double staining channels was quantified by GraphPad Prism using Pearson’s correlation coefficient (Pearson’s r), ranging between 1 and −1.

### Proximity Ligation Assay (PLA)

PLA was utilized to detect *in situ* protein-protein/PIP_n_ interaction or close proximity, as previously described^17, 18, 31^. After fixation and permeabilization, cells were blocked before incubation with primary antibodies as in routine IF staining. The cells were then processed for PLA (#DUO92101, MilliporeSigma) according to the manufacturer’s instruction and previous publications^17, 18, 31^. Post-PLA slides were further processed for immunofluorescent staining against the nuclear membrane marker (Lamin A/C, #8617, Cell Signaling). After washing, the slides were processed and imaged as indicated in the IF section above. The Leica SP8 confocal microscope detected PLA signals as discrete punctate foci and provided the intracellular localization of the complex. ImageJ was used to quantify the nuclear PLA foci.

### Crystal Violet Cell Viability Assay

In 96-well plates, 5×10^4^ cells/well of MDA-MB-231, MDA-MB-231^Cas9^, BT549, SUM159, or SUM1315 cells were transfected with control siRNAs or siRNAs targeting p53, PITPα, PITPβ, PITPNC1, or the combination of PITPα and PITPβ as indicated for 48 h. For the KD rescue experiments, MDA-MB-231 cells were transfected with control siRNAs or siRNAs targeting 3’UTR of PITPα and PITPβ for 24 h. Next, the cells were transfected with HA-tagged wild-type PITPα or its PI-binding deficient T59D mutant PITPα for another 24 h. The cells were then treated with the control vehicle or 30 μM cisplatin for 24 h, and the cells were subsequently fixed with 4% PFA and stained with 0.2% Crystal Violet. After washing, the plates were air-dried at room temperature and imaged using the EVOS M5000 Imaging System (ThermoFisher Scientific). The dye was extracted from the cells with 10% acetic acid and quantified by measuring the absorbance at 570 nm using a Synergy HTX Multi-Mode Microplate reader.

### Caspase 3 Activity Assay

In 6-well plates, 1×10^6^ cells/well of MDA-MB-231, BT549, SUM159, or SUM1315 were transfected with control siRNAs or siRNAs targeting p53, PITPα, PITPβ, PITPNC1, or the combination of PITPα and PITPβ as indicated for 48 h. For the KD rescue experiments, MDA-MB-231 cells were transfected with control siRNAs or siRNAs targeting 3’UTR of PITPα and PITPβ for 24 h. Then, the cells were transfected with either HA-tagged wild-type PITPα or its PI-binding deficient T59D mutant PITPα for another 24 h, followed by treatment with a control vehicle or 30 μM cisplatin for 24 h. Next, the cells were lysed and processed for a Caspase 3 activity assay following the manufacturer’s instructions (#E-13183, Thermo Fisher Scientific). The fluorescence was read at excitation/emission 342/441 nm using a Synergy HTX Multi-Mode Microplate reader.

### Microscale Thermophoresis (MST) Assay

The microscale thermophoresis (MST) assay was used to calculate the binding affinities of purified recombinant proteins (p53 (#81091, Active Motif), PITPα (#ab101666, Abcam), PITPβ (#LS-C794864, LSBio), PITPNC1 (#LS-G33119-10, LSBio) PI4KIIα (lab-made)) and other contributing factors *in vitro* as previously described^64^. Following the manufacturer’s instruction, the target protein was fluorescently labeled using a Monolith Protein Labeling kit (RED-NHS 2nd Generation MO-L011, Nano Temper). Then, sequential titration of unlabeled ligand proteins or PI (#840042P, Avanti), was made in a Tris-based MST buffer and mixed with an equal volume of fluorescently labelled target protein prepared at 10 nM concentration in the same MST buffer, making the final target protein at a constant concentration of 5 nM and the ligand-protein or lipid as the gradient. For PI cofactor analysis, target proteins were mixed with 1 μM PI before ligand proteins. The target–ligand mixtures were loaded into Monolith NT.115 series capillaries (MO-K022, Nano Temper), and the MST traces were measured using Monolith NT.115 pico. The binding affinity was auto-generated using MO Control v.1.6 software.

### *In vitro* binding assay

The recombinant proteins p53 (#81091, Active Motif), PITPα (#ab101666, Abcam), PITPβ (#ab101091, Abcam), and PI4KIIα (lab made) were used for *In Vitro* binding assays. The binding assay was performed in PBS by incubating a constant amount of p53 or PI4KIIα with an increasing amount of p53, PITPα/β, or PI4KIIα in the presence of 20 μl anti-p53 antibody-conjugated agarose (#sc-126AC, Santa Cruz). After incubating overnight at 4°C, unbound proteins were removed by washing three times with PBST, and the protein complex was analyzed by immunoblotting.

### Nuclear Fractionation

Nuclear fractionation was used to analyze compartmentalized protein expression. MDA-MB-231 cells were processed using the NE-PER nuclear and cytoplasmic extraction reagents (#78833, Thermo) according to the manufacturer’s instructions. Cells were manually removed from their culture dishes into suspension, and the cytosolic extraction reagent was used to isolate cytosolic fractions. Then, the nuclear extraction reagent was used to produce the nuclear fraction.

### Lenti-CRISPR Gene Editing

CRISPR-Cas9 genome editing was used to knock out the gene encoding PITPβ in MDA-MB-231 cells. First, a Cas9 expressing stable cell line was generated by transducing cells with viral particles (#LVCAS9BST, Sigma Aldrich) at the multiplicity of infection (MOI) of 0.5. Transduction was performed in complete media containing 8µg/mL polybrene. After 24 h incubation, Cas9-expressing cells were selected with 10µg/mL blasticidin (#DSB12150; Dot Scientific) for 1 week, and expression was confirmed. To generate PITPβ KO cells, viral particles containing the guide RNA sequence for PITPβ (5’-CAATCATCCTCACGAATGC-3’) and negative control (5’-CGCGATAGCGCGAATATATT-3’) were purchased from Sigma Aldrich. Sequences were cloned into LV21 lentiviral vectors (Sigma Aldrich). Cas9-expressing cells were then transduced with viral particles at a MOI of 0.5 and incubated for 24 h. Transduction was performed in the presence of 8µg/mL polybrene in complete media. Media was removed 24 h after viral transduction and 1500 µg/mL geneticin (#10131035; Sigma Aldrich) was added for selection. After 1 week of selection in antibiotics, single cells were isolated in the 96-well tissue culture dish by BD FACS AriaII.

### Transwell Invasion Assay

The bottom polycarbonate filter surface of a 6.5 mm diameter insert with 8 μm pores in a 24-well plate (#3422, Corning) was coated with 10 μg/ml of Laminin (#CC095, Millipore Sigma) diluted in PBS for 3 h at 37°C. MDA-MB-231 cells were transiently transfected with control siRNAs or siRNAs targeting the 3’UTR of both PITPα and PITPβ for 24 h. Then, the cells were transfected with either HA-tagged wild-type PITPα or its PI-binding deficient T59D mutant PITPα. After 24 h, cells were serum starved for an additional 24 h. Next, 5×10^4^ transfected cells were suspended in 200 μl serum-free medium containing 0.5% BSA and then were plated in the upper insert chamber in 500 μl serum-free medium with 0.5% BSA. 10 ng/ml EGF was added to the lower chamber. Cells were allowed to invade for 16 h at 37°C. Cells on the bottom of the filter were then fixed with 4% PFA and stained with 0.2% Crystal Violet. The stained cells were imaged using the EVOS M5000 Imaging System (ThermoFisher Scientific). At the end of the assay, the dye was extracted by 10% acetic acid from the cells and quantified by measuring the optical density at 570 nm using a Synergy HTX Multi-Mode Microplate reader.

### [^3^H]*myo*-inositol Metabolic Labeling

Cells were split and cultured in low serum Opti-MEM (#31985070, Thermo) supplemented with 10% dialyzed FBS with a 10,000 mw cut-off (#F0392, Sigma) and 1% Pen/strep (#15140-122, Gibco) to minimize inositol in the culture media. In 10 cm dishes, 0.5×10^6^ MDA-MB-231 cells were seeded to achieve ∼5% confluency the following day. 24 h after plating, cells were treated with either 1.322 μM (25 μCi/mL) of [^3^H]*myo*-inositol (#NET1156005MC, PerkinElmer) or unlabeled *myo*-inositol (#J60828.22, Thermo) and allowed to uptake the metabolic label for 48 h. For KD experiments, PITPα/β siRNAs were added and recovered as normal and fresh [^3^H]*myo*-inositol supplemented media was used to replace the transfection media. 24 h after siRNA transfection (72 h after original plating), cells were treated with control vehicle or 30 μM cisplatin for another 24 h. Labeled and unlabeled cells were then lysed and p53 was immunoprecipitated as described in *immunoprecipitation and* immunoblotting, before performing SDS-PAGE or CHCl_3_/MeOH extraction as indicated. For extraction, MeOH:CHCl_3_ (2:1) was added to RIPA cell lysates to form the protein precipitate. Soluble phases in MeOH and CHCl_3_ were aspirated out and the protein flake was dried using a speed vacuum. Protein samples were then resuspended and solubilized in PBS with sonication. SDS-PAGE samples were excised from the gel and each lane was either further sectioned or treated as a whole and dissolved in 30% H_2_O_2_ before liquid scintillation counting (LSC). Dissolved gel samples or resuspended protein samples from the CHCl_3_/MeOH extraction were added to LSC vials with LSC cocktail (#6013319, PerkinElmer) before being processed by PerkinElmer Tri-Carb 4910 TR liquid scintillation analyzer. Analysis and DPM calculation were automated using QuantaSmart software.

### Statistics and Reproducibility

Two-tailed unpaired *t*-tests were used for pair-wise significance, and one-way ANOVA was used for group significance with Bonferroni’s multiple comparisons test used for significance within grouped samples. We note that no power calculations were used. Sample sizes were determined based on previously published experiments where significant differences were observed^17, 18^. We used at least three independent experiments or biologically independent samples for statistical analysis.

### Resource and Data Availability

To allow others to replicate and build on this work, we will make all materials, data, and associated protocols available to readers in a public repository upon publication.

---

**Extended Data Figure 1.**
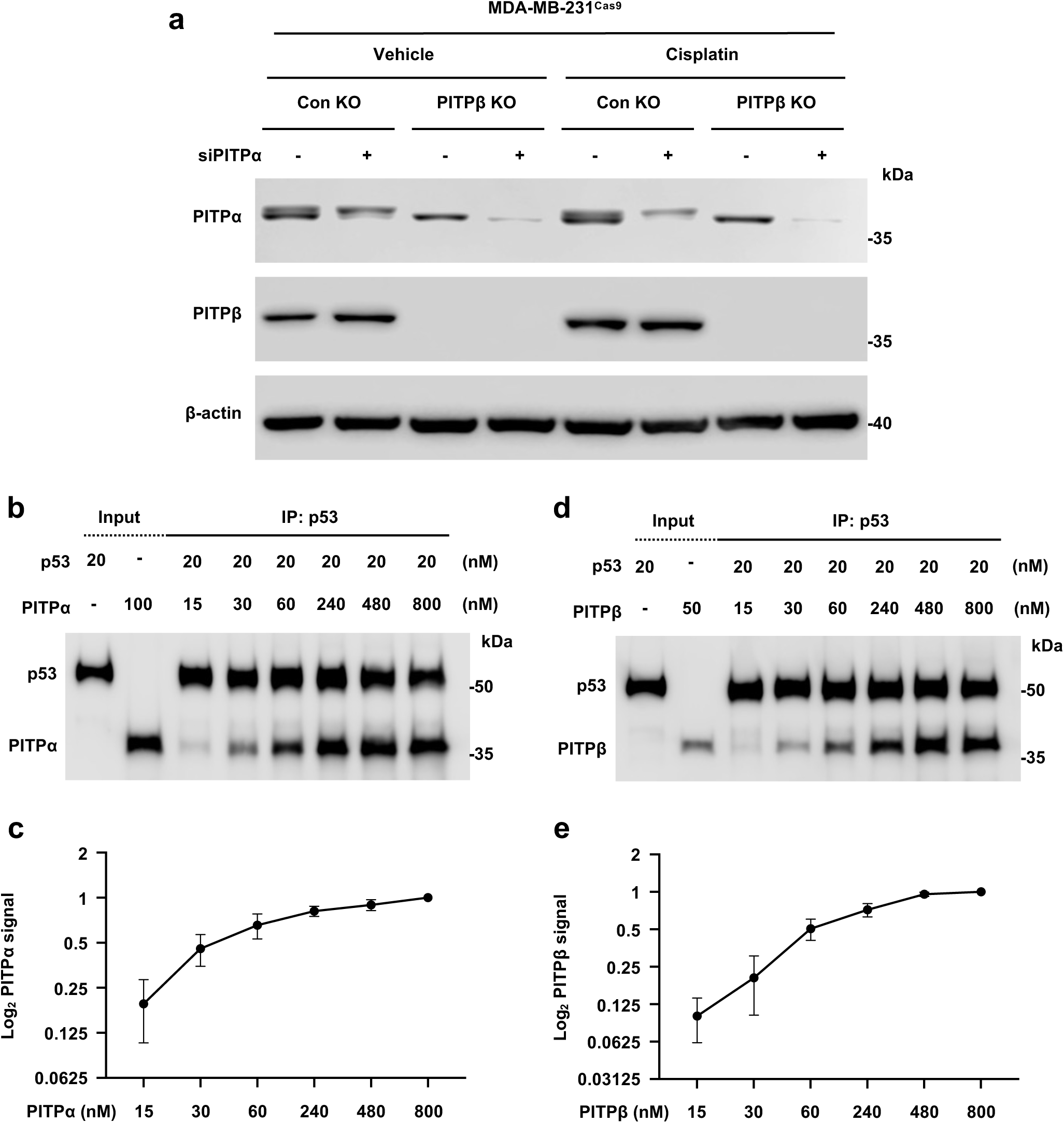
PITPα/β validation and interaction with p53. **a**, MDA-MB-231^Cas9^ cells with PITPβ KO and control non-targeted KO were transfected with control siRNAs or siRNAs against PITPα. After 48 h, cells were processed for WB to analyze PITPα and PITPβ expression. n=3 independent experiments. **b-e**, *In vitro* binding of recombinant p53 and PITPα (**b-c**) and p53 and PITPβ (**d-e**). Anti-p53 antibody-conjugated agarose was incubated with constant p53 and increasing PITPα or PITPβ protein. p53 was then IPed and analyzed by WB and quantified by ImageJ for p53-bound PITPα (**c**) and p53-bound PITPβ (**e**). n=3 independent experiments. For all graphs, data are presented as the mean ± SD.

**Extended Data Figure 2.**
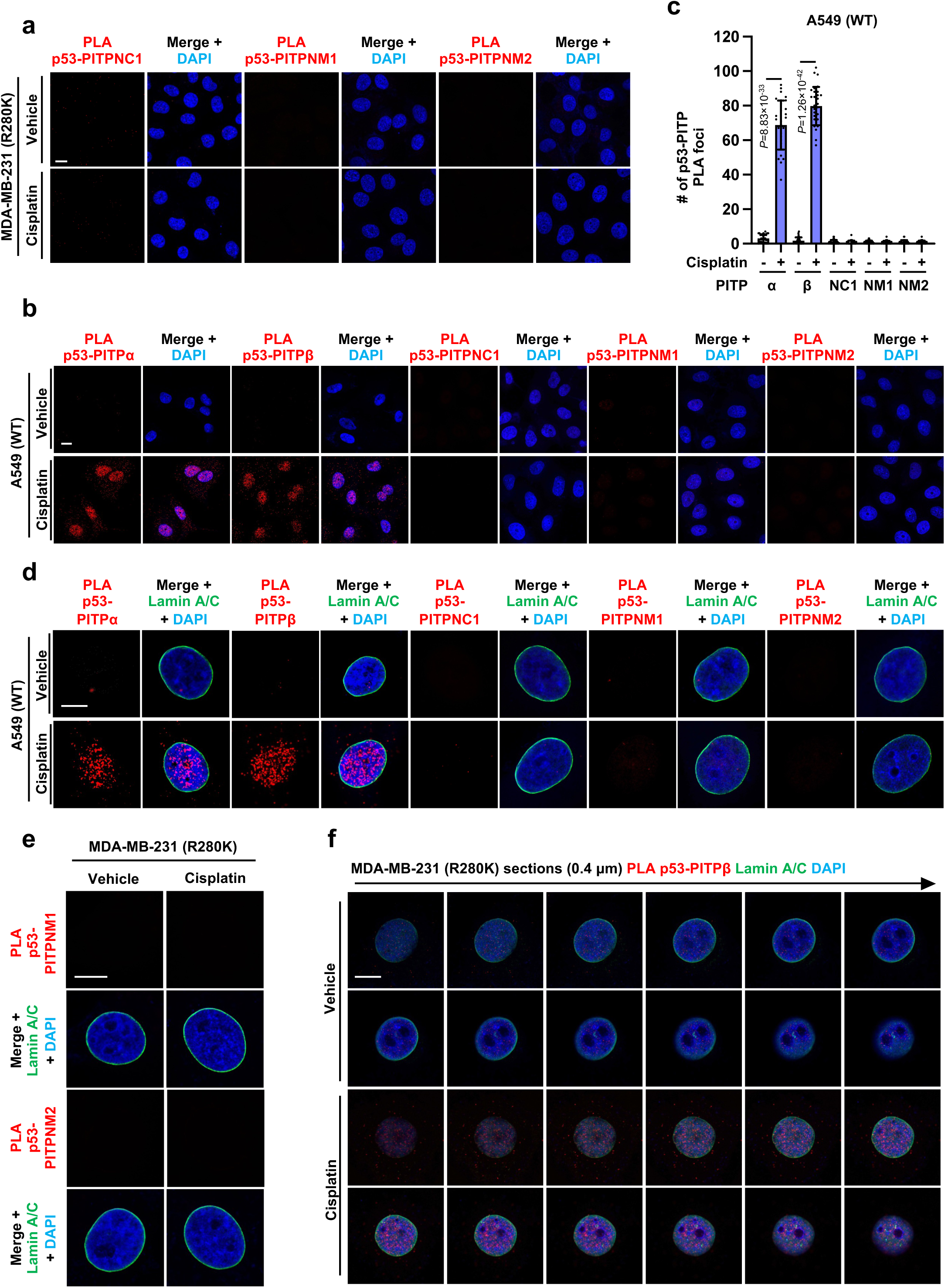
p53 interacts with class I PITPs in the nucleus in response to stress. **a**, PLA of p53-PITPα/PITPβ/PITPNC1/PITPNM1/PITPNM2 in MDA-MB-231 cells treated with vehicle or 30 µM cisplatin for 24 h. n=3, 10 cells from each independent experiment. *p* value denotes two-sided paired t-test. See expanded images in Fig. 1j and quantification in Fig. 1k. **b-c**, PLA of p53-PITPα/PITPβ/PITPNC1/PITPNM1/PITPNM2 in A549 cells treated with vehicle or 30 µM cisplatin for 24 h. The nuclear p53-PITP foci were quantified (**c**). n=3, 10 cells from each independent experiment. *p* value denotes two-sided paired t-test. **d**, PLA of p53-PITPα/PITPβ/PITPNC1/PITPNM1/PITPNM2 overlaid with the nuclear envelope marker Lamin A/C in A549 cells treated with vehicle or 30 µM cisplatin for 24 h. The nuclei were counterstained by DAPI. n=3 independent experiments. **e**, PLA of p53-PITPα/PITPβ/PITPNC1/PITPNM1/PITPNM2 overlaid with the nuclear envelope marker Lamin A/C in MDA-MB-231 cells treated with vehicle or 30 µM cisplatin for 24 h. The nuclei were counterstained by DAPI. n=3 independent experiments. See expanded images in Fig. 1l. **f**, 3D sections of p53-PITPβ PLA foci overlaid with Lamin A/C in MDA-MB-231 treated with vehicle or 30 µM cisplatin for 24 h. The nuclei were counterstained by DAPI. Each frame of the 3D sections was over a 0.2 µm thickness. For all graphs, data are presented as the mean ± SD. Scale bar, 5 µm.

**Extended Data Figure 3.**
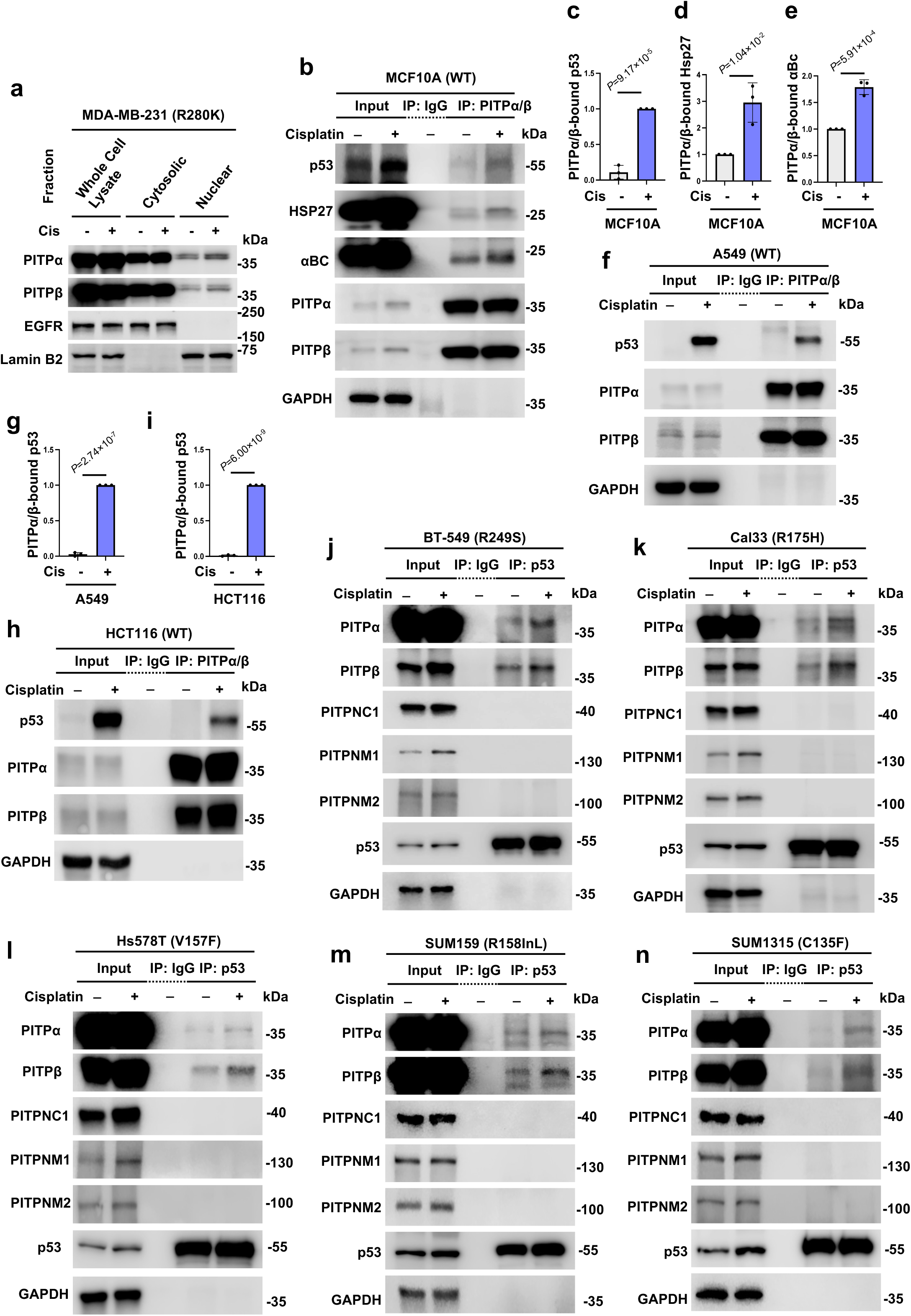
p53 interacts with class I PITPs in multiple cell lines. **a,** MDA-MB-231 cells were treated with vehicle or cisplatin for 24 h. Cells were then fractionated using a detergent-based extraction to produce lysates from cytosolic and nuclear fractions and analyzed via WB. n=3 independent experiments. **b-e,** Co-IP of p53/Hsp27/αB-crystallin (αBC) with the shared epitope of PITPα/β in MCF10A cells treated with vehicle or 30 µM cisplatin for 24 h. p53 immunoprecipitated (IPed) by PITPα/β (**c**), Hsp27 IPed by PITPα/β (**d**), and αBC IPed by PITPα/β (**e**) were analyzed by WB. n=3 independent experiments. **f-g,** Co-IP of p53 with the shared epitope of PITPα/β in A549 cells treated with vehicle or 30 µM cisplatin for 24 h. p53 IPed by PITPα/β was analyzed by WB (**g**). n=3 independent experiments. **h-i,** Co-IP of p53 with the shared epitope of PITPα/β in HCT116 cells treated with vehicle or 30 µM cisplatin for 24 h. p53 IPed by PITPα/β was analyzed by WB (**i**). n=3 independent experiments. **j-n,** Co-IP of PITPα/PITPβ with p53 in BT549, Cal33, HS578T, SUM159, SUM1315 cells treated with vehicle or 30 µM cisplatin for 24 h. PITPα/PITPβ IPed by p53 and PITPNC1/PITPNM1/PITPNM2 were analyzed by WB. n=3 independent experiments. For all panels, data are represented as mean ± SD, and the *p* value denotes a two-sided paired t-test.

**Extended Data Figure 4.**
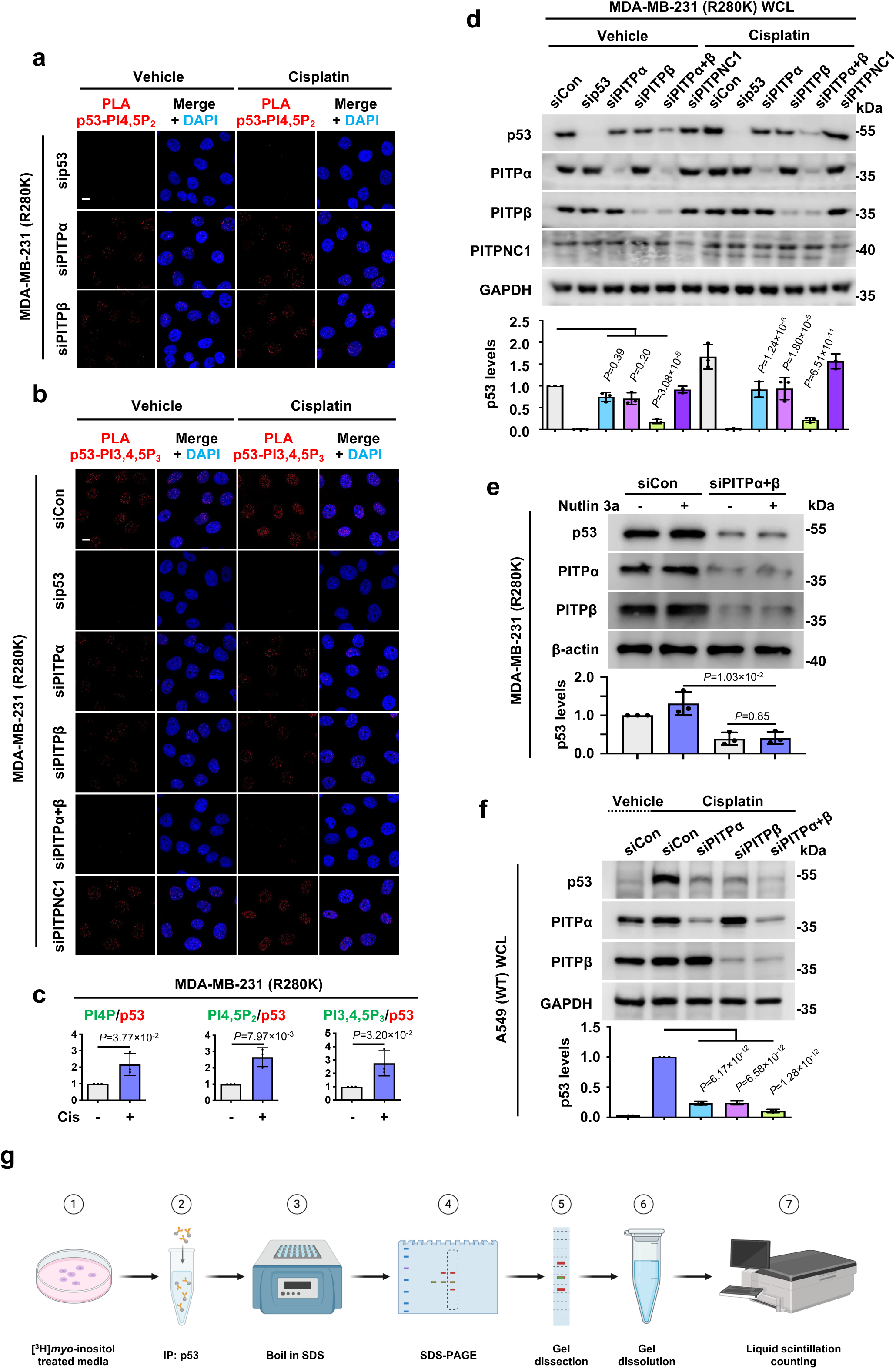
PITPα/β regulate p53-PIP_n_ complexes and p53 levels. **a-b,** MDA-MB-231 cells were transfected with control siRNAs or siRNAs against p53, PITPα, PITPβ, PITPNC1, or both PITPα and PITPβ. After 24 h, cells were treated with 30 µM cisplatin or vehicle for 24 h before being processed for PLA to detect p53-PI4,5P_2_ (**a**) and p53-PI3,4,5P_3_ (**b**) complexes. n=3 independent experiments. See KD validation in Extended Data Fig. 4d, expanded images in Fig. 2a, and quantification in Fig. 2b,c. **c,** MDA-MB-231 cells were treated with 30 µM cisplatin or vehicle for 24 h before being processed for IP against p53. p53-bound PI4P/PI4,5P_2_/PI3,4,5P_3_ were examined by fluorescent WB in Fig. 2g before being analyzed by imageJ. n=3 independent experiments. *p* value denotes two-sided paired t-test. **d**, MDA-MB-231 cells were transfected with control siRNAs or siRNAs against p53, PITPα, PITPβ, PITPNC1, or both PITPα and PITPβ. After 24 h, cells were treated with 30 µM cisplatin or vehicle for 24 h before being processed for WB to validate the KD. The p53 level was quantified by ImageJ. n=3 independent experiments. *p* value denotes ANOVA with Bonferroni’s multiple comparisons test. **e**, MDA-MB-231 cells were transfected with control siRNAs or siRNAs against both PITPα and PITPβ. After 24 h, cells were treated with 10 µM Nutlin 3a or vehicle for 24 h before being processed for WB and analyzed using ImageJ. *p* value denotes two-sided paired t-test. n=3 independent experiments. **f**, A549 cells were transfected with control siRNAs or siRNAs against PITPα, PITPβ, or both PITPα and PITPβ. After 24 h, cells were treated with 30 µM cisplatin or vehicle for 24 h before being processed for WB. The p53 level was quantified by ImageJ. n=3 independent experiments. *p* value denotes ANOVA with Bonferroni’s multiple comparisons test. **g**, A model of the [^3^H]-inositol lableing workflow used to detect p53-PIP_n_ complexes. See related data in Fig. 3a-b. For all panels, data are represented as mean ± SD. Scale bar, 5 μm.

**Extended Data Figure 5.**
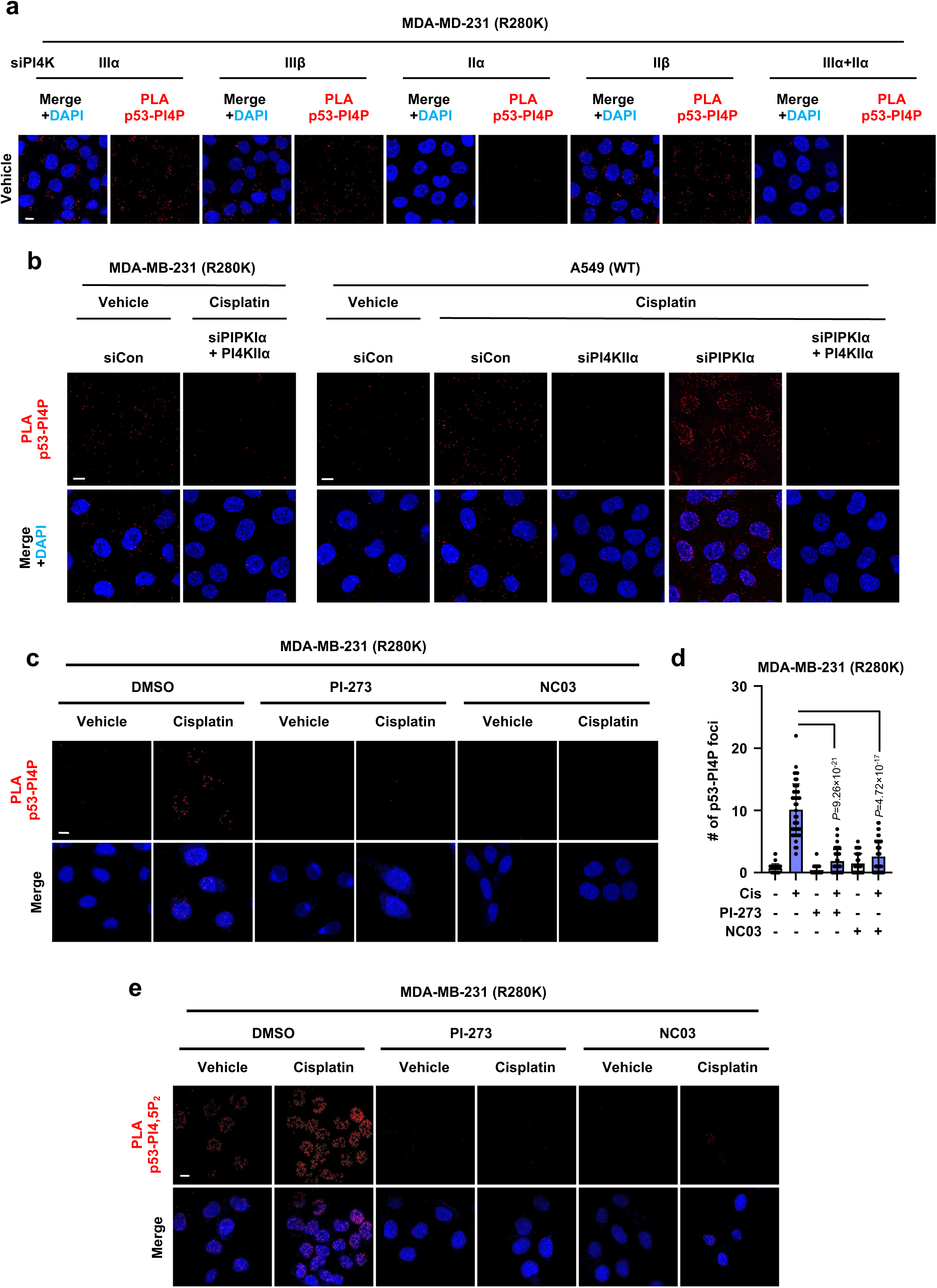
PI4KIIα activity is required to generate p53-PI4P. **a**, MDA-MB-231 cells were transfected with control siRNAs or siRNAs against PI4KIIIα, PI4KIIIβ, PI4KIIα, PI4KIIβ or both PI4KIIα and PI4KIIIα. After 24 h, cells were treated with 30 µM cisplatin or vehicle for 24 h before being processed for PLA to detect p53-PI4P complexes (**b**). n=3 independent experiment. See expanded images in Fig. 4a and quantification in Fig. 4b. **b**, MDA-MB-231 and A549 cells were transfected with control siRNAs or siRNAs against PI4KIIIα, PIPKIα or both PI4KIIα and PIPKIα. After 24 h, cells were treated with 30 µM cisplatin or vehicle for 24 h before being processed for PLA to detect p53-PI4P complexes. n=3 independent experiment. See KD confirmation in Fig. 4d, expanded images in Fig. 4e, and quantification in Fig. 4f. **c-d**, MDA-MB-231 cells were treated with vehicle or cisplatin in combination with DMSO as a control, PI-273, or NC03 for 24 h before being processed for PLA to detect p53-PI4P complexes and analyzed using ImageJ (**d**). *p* value denotes two-sided paired t-test. n=3, 15 cells from each independent experiment. **e**, MDA-MB-231 cells were treated with vehicle or cisplatin in combination with DMSO as a control, PI-273, or NC03 for 24 h before being processed for PLA to detect p53-PI4,5P_2_ complexes. n=3 independent experiments. See quantification in Fig. 4g. For all panels, scale bar, 5 μm.

**Extended Data Figure 6.**
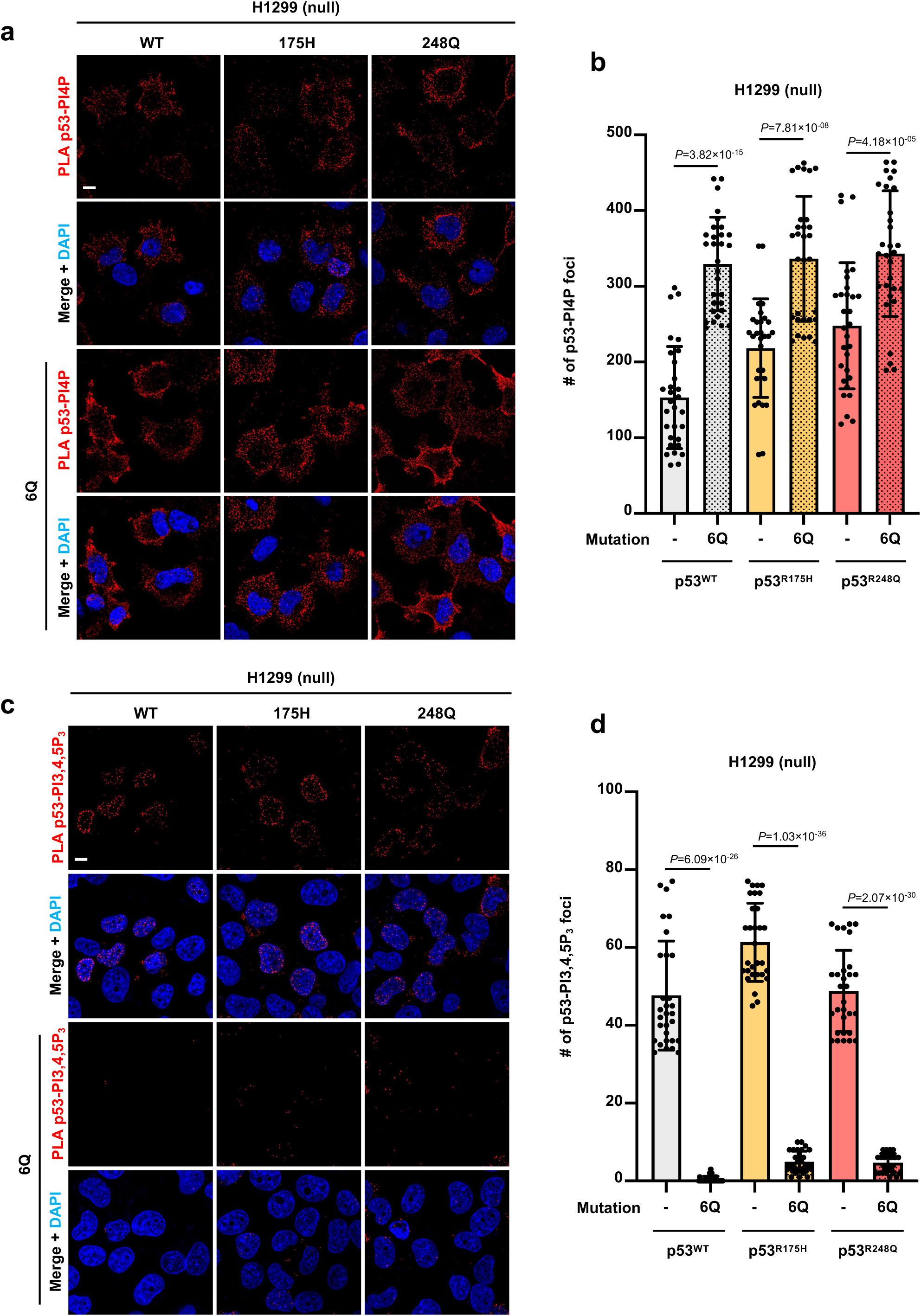
p53^6Q^ mutants still form p53-PI4P complexes. **a-b,** H1299 cells transfected with constructs expressing p53 WT, WT6Q, 175H, 175H6Q, 248Q and 248Q6Q were treated with 30 µM cisplatin for 24 h. Then, the cells were processed for PLA to detect p53-PI4P complexes (**b**). Nuclei were counterstained by DAPI. *p* value denotes two-sided paired t-test. n=3 independent experiments. **c-d,** H1299 cells transfected with constructs expressing p53 WT, WT6Q, 175H, 175H6Q, 248Q, and 248Q6Q were treated with 30 µM cisplatin for 24 h. Then, the cells were processed for PLA to detect p53-PI3,4,5P_3_ complexes (**d**). Nuclei were counterstained by DAPI. *p* value denotes two-sided paired t-test. n=3 independent experiments. For all panels, data are represented as mean ± SD. Scale bar, 5 μm.

**Extended Data Figure 7.**
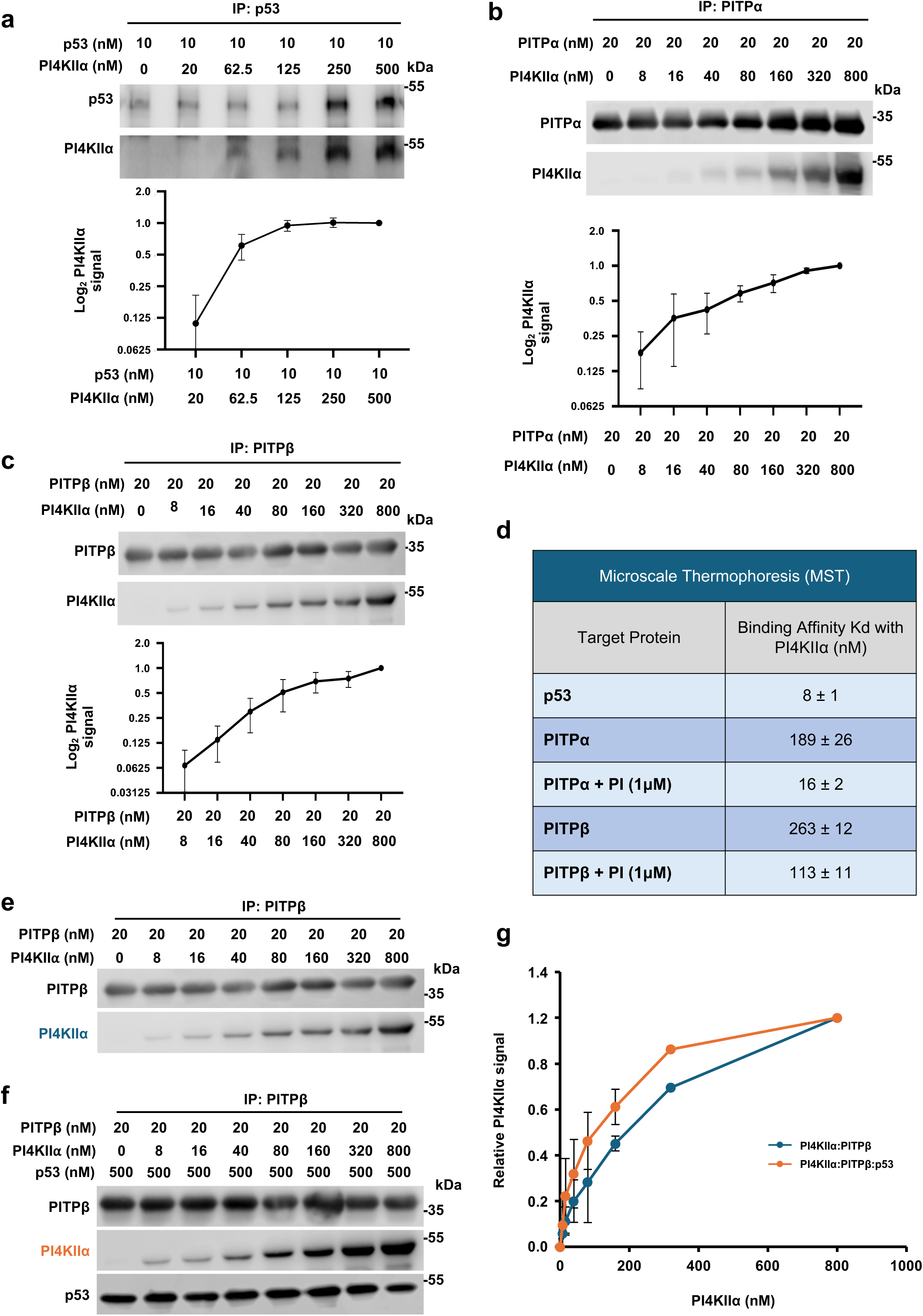
PΙ4ΚΙΙα interacts with PITPα/β and p53 cooperatively. **a-c**, *In vitro* binding of recombinant PI4KIIα and p53 (**a**), PI4KIIα and PITPα (**b**), and PI4KIIα and PITPβ (**c**). p53, PITPα, or PITPβ immobilized on the indicated antibody-conjugated agarose was incubated with PI4KIIα protein. p53, PITPα or PITPβ was IPed and p53-bound PI4KIIα (**a**), PITPα-bound PI4KIIα (**b**) and PITPβ-bound PI4KIIα (**c**) was analyzed by WB and quantified by ImageJ. n=3 independent experiments. **d**, The interaction of recombinant fluorescently labelled PI4KIIα with other components was quantitated by MST assay. A constant concentration of fluorescently labelled PI4KIIα (5 nM) was incubated with increasing concentrations of non-labelled ligand with or without the addition of 1 μM PI and analyzed using a Monolith NT.115 pico, and the binding affinity was autogenerated by MO. Control v.1.6 software. The binding affinities determined by MST are shown as indicated K_d_ values. n=3 independent experiments. **e-g**, *In vitro* binding of recombinant PITPβ and PI4KIIα with and without p53. Anti-PITPβ antibody-conjugated agarose was incubated with constant PITPβ and increasing PI4KIIα protein in the absence (**e**) or presence (**f**) of p53. PITPβ was then IPed, analyzed by WB, and quantified by ImageJ for PITPβ-bound PI4KIIα. n=3 independent experiments. For all panels, data are presented as the mean ± SD.

**Extended Data Figure 8.**
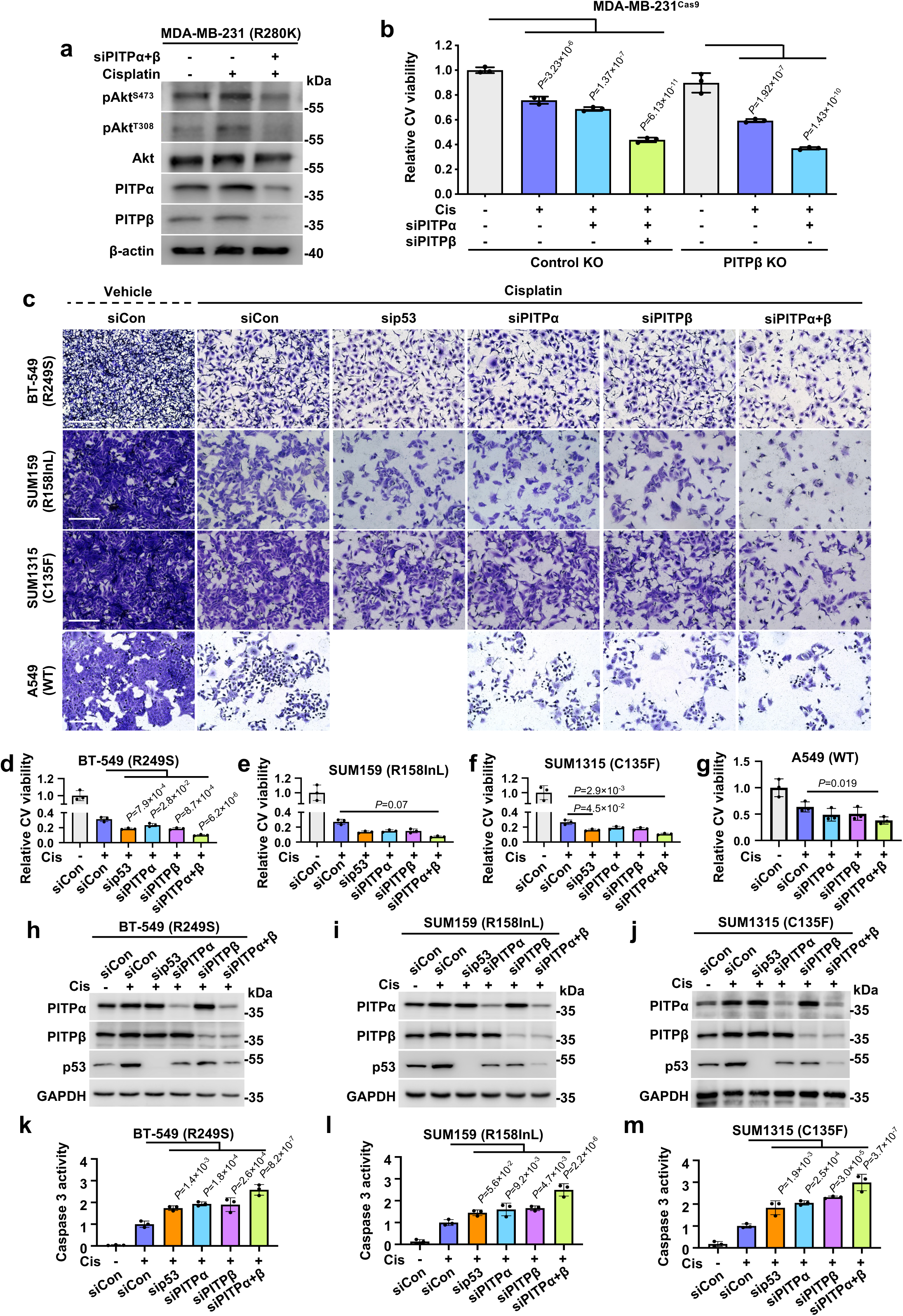
PITPα/β regulate Akt activation and chemoresistance. **a**, MDA-MB-231 cells were transfected with control siRNAs or siRNAs against both PITPα and PITPβ. After 24 h, cells were treated with 30 µM cisplatin or vehicle for 24 h before being processed for WB. n=3 independent experiments. **b**, MDA-MB-231^Cas9^ cells with PITPβ KO and control non-targeted KO were transfected with control siRNAs or siRNAs against PITPα, PITPβ, or both PITPα and PITPβ. After 24 h, cells were treated with 30 µM cisplatin or vehicle for 24 h before being processed for Crystal Violet viability assay. Viability was quantified based on the extracted dye using a plate reader. *p* value denotes ANOVA with Bonferroni’s multiple comparisons test. n=3 independent experiments. See representative images in Fig. 5h. **c-j**, BT-549, SUM159, and SUM1315 cells were transfected with control siRNAs or siRNAs against p53, PITPα, PITPβ, or both PITPα and PITPβ. A549 cells were transfected with control siRNAs or siRNAs against PITPα, PITPβ, or both PITPα and PITPβ. After 24 h, cells were treated with 30 µM cisplatin or vehicle for 24 h before being processed for Crystal Violet viability assay. The cells were imaged by an EVOS M5000 microscope (**c**) and quantified based on the extracted dye using a plate reader (**d-f**). The KD was validated by WB (**g-i**). *p* value denotes ANOVA with Bonferroni’s multiple comparisons test. n=3 independent experiments. **k-m**, BT-549, SUM159, and SUM1315 cells were transfected with control siRNAs or siRNAs against p53, PITPα, PITPβ, or both PITPα and PITPβ. After 24 h, cells were treated with 30 µM cisplatin or vehicle for 24 h before being processed for a caspase 3 activity assay. *p* value denotes ANOVA with Bonferroni’s multiple comparisons test. n=3 independent experiments. For all graphs, data are presented as the mean ± SD. Scale bar, 300 µm.

**Extended Data Figure 9.**
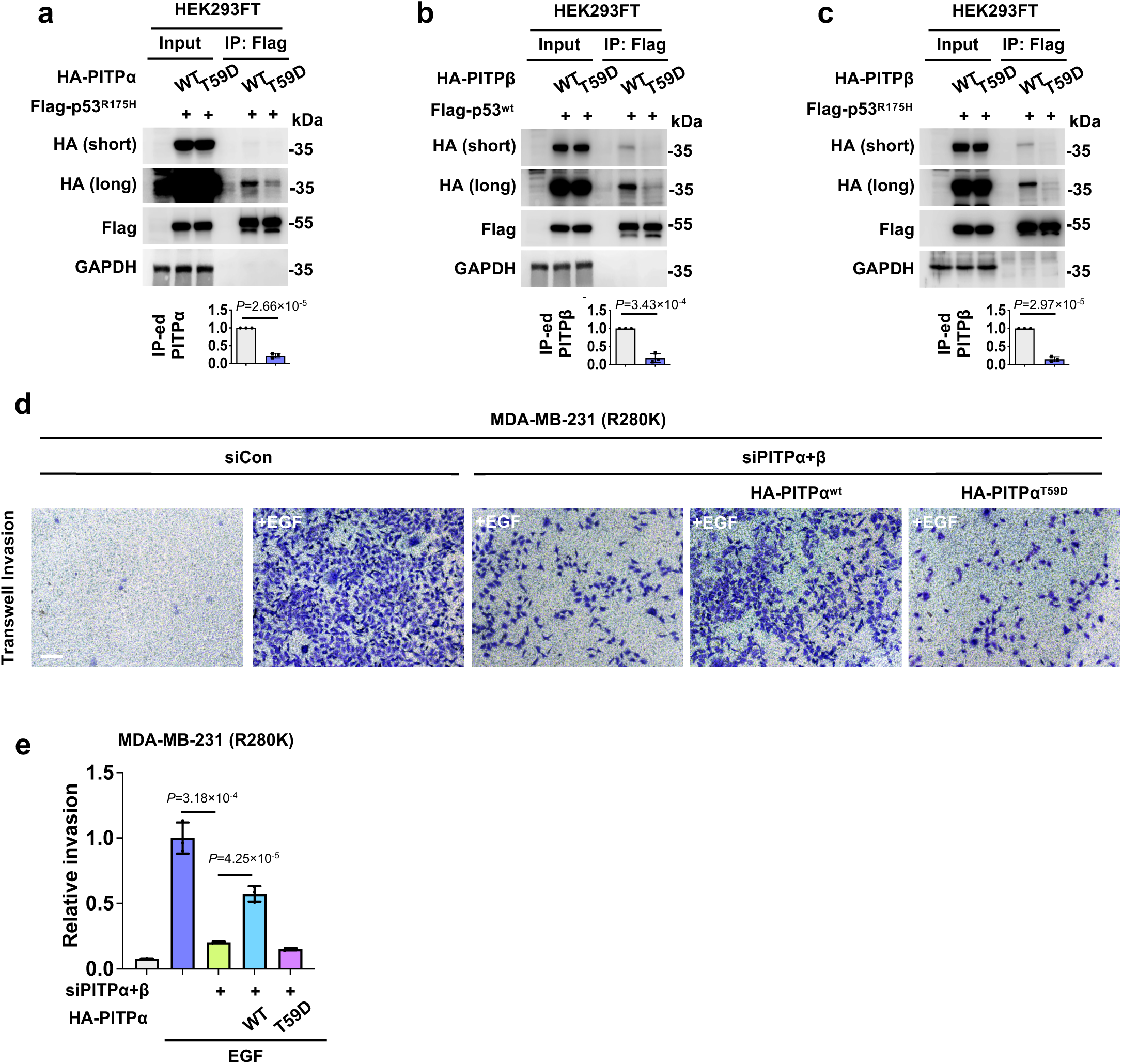
PITPs require PI binding to function with p53. **a**, HEK293FT cells were co-transfected with either HA-tagged wild-type PITPα or PI binding-defective mutant T59D PITPα together with Flag-tagged mutant p53^R175H^. After 48 h, cells were processed for IP against Flag-tag and analyzed by WB. *p* value denotes two-sided paired t-test. n=3 independent experiments. **b**, HEK293FT cells were co-transfected with either HA-tagged wild-type PITPβ or PI binding-defective mutant T59D PITPβ together with Flag-tagged p53^wt^. After 48 h, cells were processed for IP against Flag-tag and analyzed by WB. *p* value denotes two-sided paired t-test. n=3 independent experiments. **c**, HEK293FT cells were co-transfected with either HA-tagged wild-type PITPβ or PI binding-defective mutant T59D PITPβ together with Flag-tagged mutant p53^R175H^. After 48 h, cells were processed for IP against Flag-tag and analyzed by WB. *p* value denotes two-sided paired t-test. n=3 independent experiments. **d-e**, MDA-MB-231 cells were transfected with control siRNAs or siRNAs against the 3’UTR of both PITPα and PITPβ for 24 h. The cells were then transfected with either HA-tagged wild-type PITPα or PI binding-defective mutant T59D PITPα. After 24 h, cells were serum starved for an additional 24 h and then scored for invasion through Laminin-coated transwell inserts with 8 µm pores using 10 ng/ml EGF as a chemoattractant for 16 h. The invading cells at the insert bottom were stained with Crystal Violet, imaged (**d**), and quantified (**e**) based on the extracted dye using a plate reader. n=3 independent experiments. *p* value denotes two-sided paired t-test. Scale bar, 300 µm. For all graphs, data are presented as the mean ± SD.

## Notes

### Competing Interest Statement

The authors have declared no competing interest.

### Summary of Updates

This revision is largely wording and methodology updates. Some data has been moved to a companion manuscript that will be uploaded to a preprint sever as well. As a result, some text in the manuscript, figure legends, and methods has been updated accordingly.

